# Multiple cell types support productive infection and dynamic translocation of infectious Ebola virus to the surface of human skin

**DOI:** 10.1101/2024.07.19.604135

**Authors:** Kelly N. Messingham, Paige T. Richards, Anthony Fleck, Radhika A. Patel, Marija Djurkovic, Jonah Elliff, Samuel Connell, Tyler P. Crowe, J. Pablo Munoz Gonzalez, Francoise Gourronc, Jacob A. Dillard, Robert A. Davey, Al Klingelhutz, Olena Shtanko, Wendy Maury

**Author notes:** Corresponding author: 4-370 Bowen Science Building., 51 Newton Rd., Iowa City, IA 52242 USA, 1-319 335 8021.

## Abstract

Ebola virus (EBOV) within the *Filoviridae* family causes severe human disease. At late stages of infection, EBOV virions are found on the surface of patients’ skin; however, the permissive cell types within the skin and how infectious virus translocates to the apical skin surfaces is not known. Here, we describe a human transwell skin explant culture model and show that EBOV infection of human skin tissues via the basal media results in a time- and dose-dependent increase in infectious virus in dermal and epidermal tissue. Infectious virus was detected on the apical epidermal surface within 3 days, indicating that the virus propagates within and traffics through the tissue. In the dermis, EBOV-infected cells were of myeloid, endothelial and fibroblast origins, whereas keratinocytes harbored virus in the epidermis. Complementary studies showed that both purified skin fibroblasts and keratinocytes supported EBOV infection ex vivo and that both cell types required the phosphatidylserine receptor, Axl, and the endosomal protein, NPC1, for virus entry. Our experimental platform identified new susceptible cell types and demonstrated dynamic trafficking of EBOV virions that resulted in infectious virus on the skin surface; findings that may explain person-to-person transmission via skin contact.

**Teaser:** Using a human skin explant model, these studies identify and characterize skin cell populations that support Ebola virus infection.

## Introduction

Ebola virus (EBOV) is an enveloped, negative-sense RNA virus within the *Filoviridae* family responsible for sporadic outbreaks in Central and Western Africa. The 2013-2016 West African epidemic is the largest to date, resulting in ∼28,000 infections and ∼11,000 deaths (*1–3*). More recently, several regions of the Democratic Republic of the Congo have had episodic outbreaks that persisted for weeks to months (*4*). The characteristic signs of EBOV include hemorrhagic fever, gastrointestinal symptoms, and multiple organ dysfunction syndrome (*5*). Cutaneous manifestations, including a maculopapular rash and petechiae, can be associated with EBOV infection of humans and non-human primates (NHPs) (*6*).

EBOV is spread by direct contact with an infected individual or their body fluids (*5*), with mucosal transmission considered to be an important route (*7*). Anecdotal evidence from events during the 2013-2016 West African EBOV epidemic suggests that the virus is present on the skin of infected individuals at late times during infection and on patients who have succumbed to infection (*8*). This implicates skin contact as a potentially important source of person-to-person transmission. Specifically, the 2013 spread of EBOV from Guinea into Sierra Leone was traced back to traditional funeral practices, which included participants preparing and touching the body of the deceased (*8–10*). It has also been established that EBOV RNA and infectious virus are on the skin surface at late stages of infection (*7*) and EBOV antigen and/or RNA is present in skin samples from humans and experimentally infected animals (*6, 11–14*). However, the mechanism(s) by which the virus travels to the skin surface, and the skin cells responsible for supporting infection, are not fully understood.

The lack of robust and reproducible models to study EBOV skin infection is a major barrier to defining the role of the skin in viral load and transmission. In this report, we describe an *ex vivo* model of EBOV infection of human skin explants that robustly supports infection. We identify permissive dermal and epidermal cell subsets that support EBOV infection and demonstrate that EBOV spreads through the skin from the basal surface of the explant, trafficking through the dermis and epidermis to the apical surface of the skin. Our findings identify a novel route through which infectious virus traverses the skin to the epidermal surface thereby contributing to person-to-person transmission. Further, we highlight the utility of this readily available model system for antiviral studies, providing a novel highly relevant platform for testing new drugs.

## Results

### Human skin explant model of EBOV infection

EBOV antigen-positive cells have been found in the skin of infected humans, NHPs, and guinea pigs by immunohistochemical staining of fixed tissue sections (*12, 23*). Skin explants maintained at the air-liquid interface have been favored as pre-clinical models to evaluate cosmetic and therapeutic toxicity because of the long-term maintenance of tissue integrity and viability in culture (*24–26*), but have not been previously used to assess filovirus infection. To better understand EBOV interactions with skin cells, we developed an *ex vivo* infection model utilizing full-thickness explants derived from healthy adult skin (**Fig. 1a**). Briefly, replicate 3-10 mm punch biopsies were placed dermal side down on a transwell insert so the basal surface was in contact with culture media. In our hands, uninfected explants cultured with media refreshment daily or every other day resulted in limited lactate dehydrogenase (LDH) release, indicating viability is maintained (**Suppl. Fig. 1**). Virus was inoculated into the basal media and virus infection was monitored for 12-17 days (**Fig. 1a**).

**Fig. 1.**
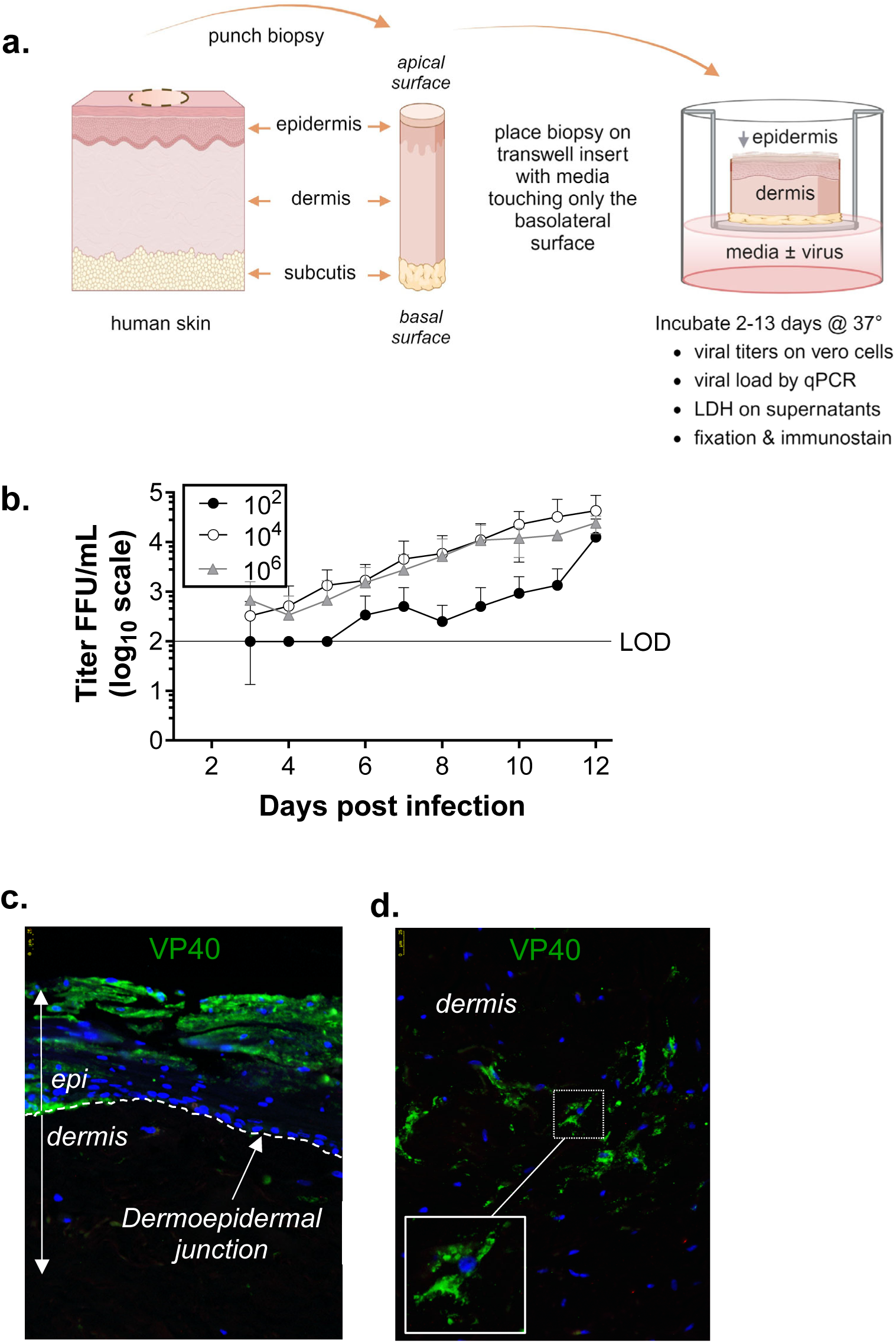
Human skin explant model of EBOV infection. **a**) Generation of skin explant cultures from human skin. Full-thickness skin punch biopsies (3-10 mm) were placed dermal side down on a transwell insert in a well containing sufficient media to maintain the explant at the air-liquid interface. Within 24 hours of culture initiation, the virus was inoculated into the culture media, and explants were incubated for up to 2 weeks with media changed every other day. Specifics regarding the virus type, dosages used, and inoculation timing are described in legends for each experiment. **b**) Infectious virus was evaluated in culture media after EBOV-GFP infection of explants over time. Media was inoculated with 10^2^, 10^4^, or 10^6^ FFU of EBOV-GFP for 24 hours and then the explants were washed and moved to a fresh insert/well. Supernatants were collected daily starting on day 3 and infectious virus titers were determined on Vero cells as previously described(*63*). Shown is one of 5 independent experiments and data is expressed as mean ± SD. **c-d**) Formalin-fixed and paraffin-embedded (FFPE) skin explants from day 12 of EBOV infection were stained with antibodies specific for EBOV GP (green) and mounted with DAPI nuclear stain. The junction of the epidermal (epi) and dermal skin layers is indicated by the dashed line. EBOV replication was observed in epidermal keratinocytes (**c**) and dermal cells (**d**) with a stellate morphology suggestive of fibroblasts (inset). Scale bar = 25 μM.

The susceptibility of fresh skin explant cultures to EBOV infection was tested by the inoculation of increasing doses (10^2^ to 10^6^ infectious units (FFU)) of EBOV-GFP and titers of infectious virus were measured in the supernatant starting on day 3. Virus titers increased in both a dose- and time-dependent manner (**Fig. 1b**). To visualize viral localization within the skin, replicate explants were formalin-fixed paraffin-embedded (FFPE) and sectioned for immunostaining. Early days after infection, EBOV-VP40 expression was confined to isolated and infrequent dermal cells, but clusters of VP40^+^ cells were evident by days 7-8 and these foci typically increased in size and frequency throughout the experiment. At later times of infection, robust viral infection was observed in both the dermis and epidermis using immunofluorescent staining (**Fig. 1c-d**), which is less sensitive than the titering assay or qRT-PCR. Together, these findings provide strong evidence that cultured skin explants serve as an excellent model for studying filovirus interactions with skin.

### Identification of dermal and epidermal cells permissive for EBOV

Human skin is a complex organ composed of a variety of structural and immune cell types. To identify dermal and epidermal cells within the explant that support EBOV infection, FFPE tissues from days 4-13 of infection were sectioned and immunostained for virus antigen (VP40, GP, or GFP) and cell lineage markers; HLA-DR (MHC class II), IBA1 (macrophages), CD31(endothelia), CD140ab (fibroblasts, smooth muscle, pericytes), CD140b (fibroblast, smooth muscle) and CK5 (keratinocytes). The specific information describing antibodies, dilutions, and staining conditions is detailed in **Table 2**.

Based on our initial studies showing that viral immunostaining was reliably evident on later days of infection, phenotypic identification of virally-infected cells was limited to days 8-13. Within the dermis, viral antigen colocalized with cells expressing HLA-DR, IBA1, CD31, and CD140ab (**Fig. 2, and Figure S2 a, b**). Viral infection of HLA-DR^+^ cells (**Fig. 2a, b**) was anticipated since myeloid cells are well-established as early and sustained cells targeted by filoviruses (*27–29*). Consistent with this, viral antigen also co-localized with dermal IBA1^+^ cells, indicating that macrophages support EBOV replication in the skin (**Fig. 2c, d**). Detection of virus in CD31^+^ cells (**Fig. 2g**) and CD140ab^+^ cells and the linear configuration of dermal CD140-positive cells suggested that components of the dermal vasculature support EBOV infection (**Fig. 2e-g**). Virus infection of dermal CD140ab^+^ spindle-shaped cells (**Fig. 2e, f and Fig. S2b**) also indicated that dermal fibroblasts are permissive. Examples of staining controls (mock infection and primary omit) are shown in **Figure S2d-i**. These findings align with studies in other tissues that report viral infection of myeloid cells, endothelial cells, and fibroblasts (*27, 28, 30, 31*).

**Fig. 2.**
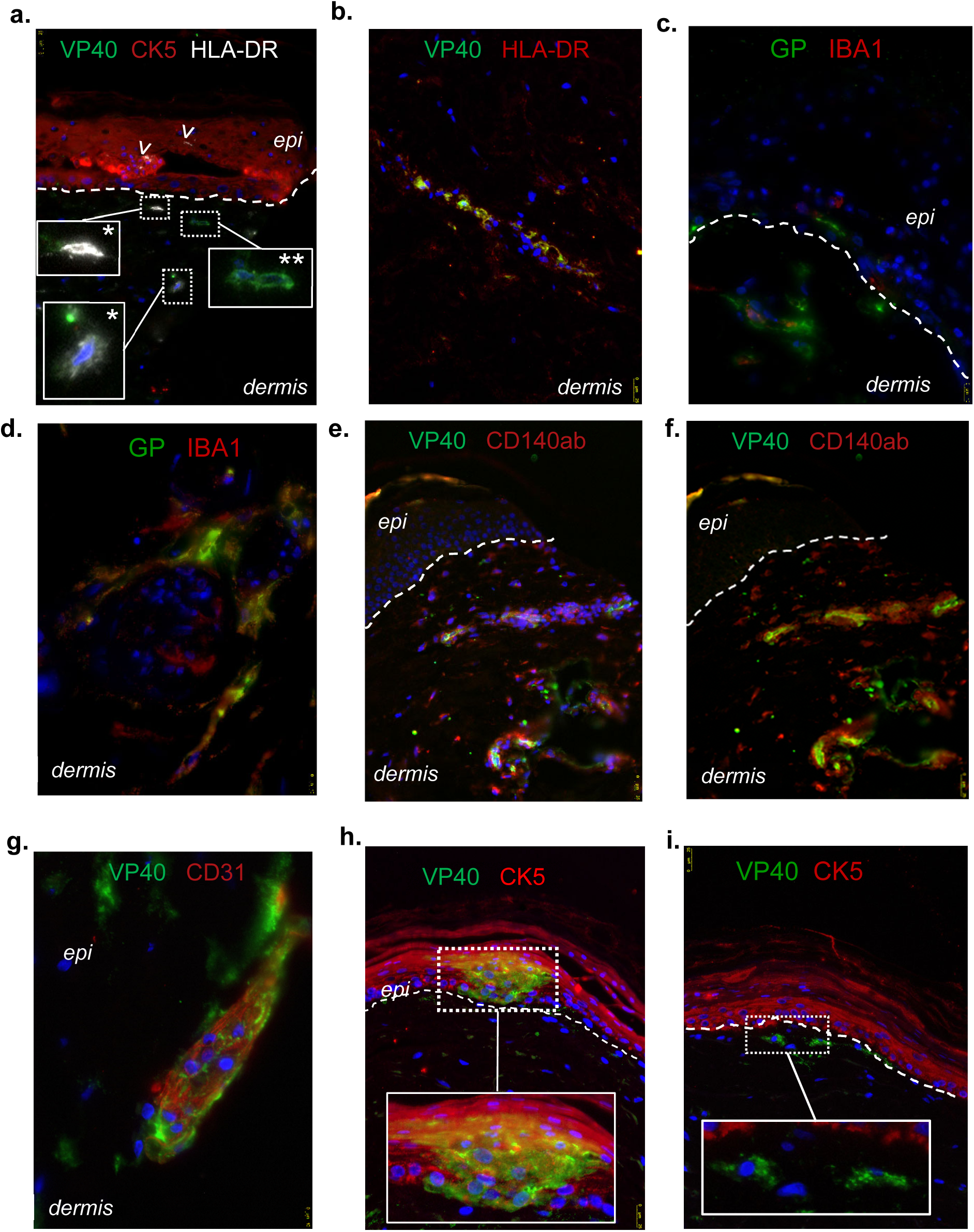
Multiple skin cell subsets support EBOV infection. Human skin explants were generated and maintained as described in Figure 1. In this experiment, explants were inoculated with 10^6^ FFU EBOV for 12 days. Sections of FFPE explants were immunostained with EBOV VP40 (green), and the indicated lineage-specific antibodies (CK5 (epidermis), HLA-DR (antigen-presenting cell), CD140ab (fibroblast/pericyte), CD31 (endothelia)) are represented in red or white and mounted with DAPI (blue). Scale bar = 10 μM in c, d, g and 25 μM in all others. The dashed line indicates the junction between the epidermis (epi) and the dermis. **a)** Viral staining was detected in dermal cells (magnified in insets) that were either HLA-DR^+^ (*) or HLA-DR-(**). Within the epidermis, HLA-DR^+^ Langerhans cells were noted (arrowheads), but viral staining was not readily apparent. **b**) Co-localization of VP40 and HLA-DR potentially along the dermal vasculature. **c, d**) virus-positive IBA1^+^ dermal macrophages **e-g**) VP40 colocalizes with markers of dermal vasculature, including CD140ab (shown with (**e**) and without (**f**) DAPI) and CD31 (**g**). Additionally, VP40 colocalizes CD140ab^+^ cells that occur singly or in small clusters in the dermis. **h)** Foci of strong VP40 staining colocalize with CK5^+^ epidermis and (**i**) in single cells in the superficial dermis. Shown are representative images from 3-5 independent experiments. Routine staining controls are shown in Supplemental Fig. 2c, d, and f.

Prior histologic studies have postulated that infectious EBOV virus reaches the skin surface by transiting through the vasculature, sweat glands, or breaks in the skin (*11*), while viral infection of epidermal cells (keratinocytes) has not been described. Thus, the colocalization of robust viral staining within the CK5^+^ epidermal keratinocytes was unexpected (**Fig. 2h, Suppl. Fig 2c, f, i, Suppl. Movie 1**). Using our transwell explant model, viral infection of dermal cell populations typically preceded the epidermis. Epidermal infection was observed at the late time points (≥day 8) and was often accompanied by virus-positive cells, such as fibroblasts, in the superficial dermis (**Fig. 2h, i**).

To determine if actively dividing basal keratinocytes were principally responsible for supporting EBOV infection in the epidermis, EBOV-infected skin sections were costained for virus antigen and MCM2, a well-established marker of dividing cells (*32, 33*). As expected, CK5^+^ basal keratinocytes were the most prominent cycling cell population (*34*) although MCM2-positive cells were evident in both the epidermis and dermis of the skin explants (**Fig. 3**). Throughout the skin, EBOV antigen was detected in both MCM2^+^ and MCM2^-^ cells, indicating that infection was not dependent upon active cell division.

**Fig. 3.**
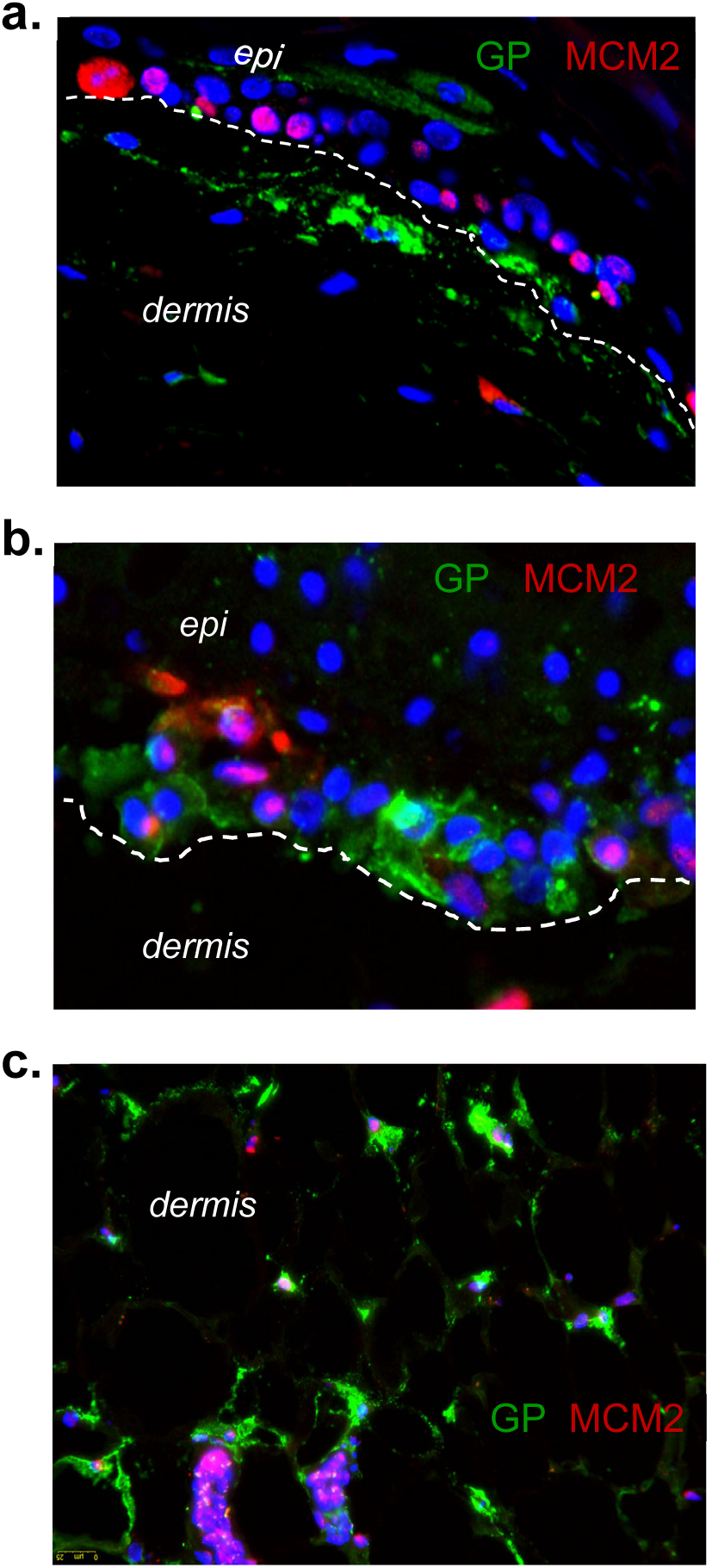
Cycling and quiescent skin cells within the explants are susceptible to EBOV infection. Human skin explants were generated and maintained as described in Fig. 1. Explants were inoculated with 10^6^ FFU EBOV and cultured for 13 days. Sections of FFPE explants were immunostained with EBOV GP (green), and MCM2 (red), a marker of cellular proliferation. Viral replication was observed in both cycling and quiescent cells in the epidermis (**a, b**) and dermis (**c**) of skin explants.

### rVSV/EBOV GP infection of skin explants as a model of EBOV infection

We next performed studies using a low containment model virus rVSV/EBOV GP that is composed of recombinant vesicular stomatitis virus (VSV) with its native G glycoprotein gene replaced with EBOV GP and GFP. In previous studies, this infectious virus was shown to have the same tropism as EBOV and has been extensively used to dissect the filoviral entry pathway (*16, 35–41*). Initial studies with foreskin explants demonstrated that input of 5 x 10^5^ iu rVSV/EBOV GP or greater resulted in infection and spread (**Suppl. Fig. 3a**), so subsequent studies with adult skin explants routinely used 10^7^ iu of the virus. Further, we found that infections in the presence of the type I interferon (IFN) inhibitor, B18R (*42*), or the Jak inhibitor, ruxolitinib (rux), enhanced infectious titers produced by these explants. This was anticipated since VSV does not encode proteins with IFN antagonistic activity and is controlled by the IFN pathway (*43*). With the utility of the rVSV/EBOV GP model virus confirmed, an extended time course of infection was performed in adult skin explants. Explants infected with rVSV/EBOV GP resulted in the detection of infectious virus in supernatants over the 13 days with TCID_50_ assay (**Fig. 4a**). The presence of the IFN inhibitor, B18R, in the media significantly enhanced virus titers at most time points. Similarly, viral loads (qRT-PCR) also increased over time and were enhanced by the addition of an IFN inhibitor (**Fig. 4b**). Explant viability as assessed by supernatant LDH levels suggested that the explants remained healthy through 12-days of rVSV/EBOV GP infection (**Suppl. Fig. 3b**).

**Fig. 4.**
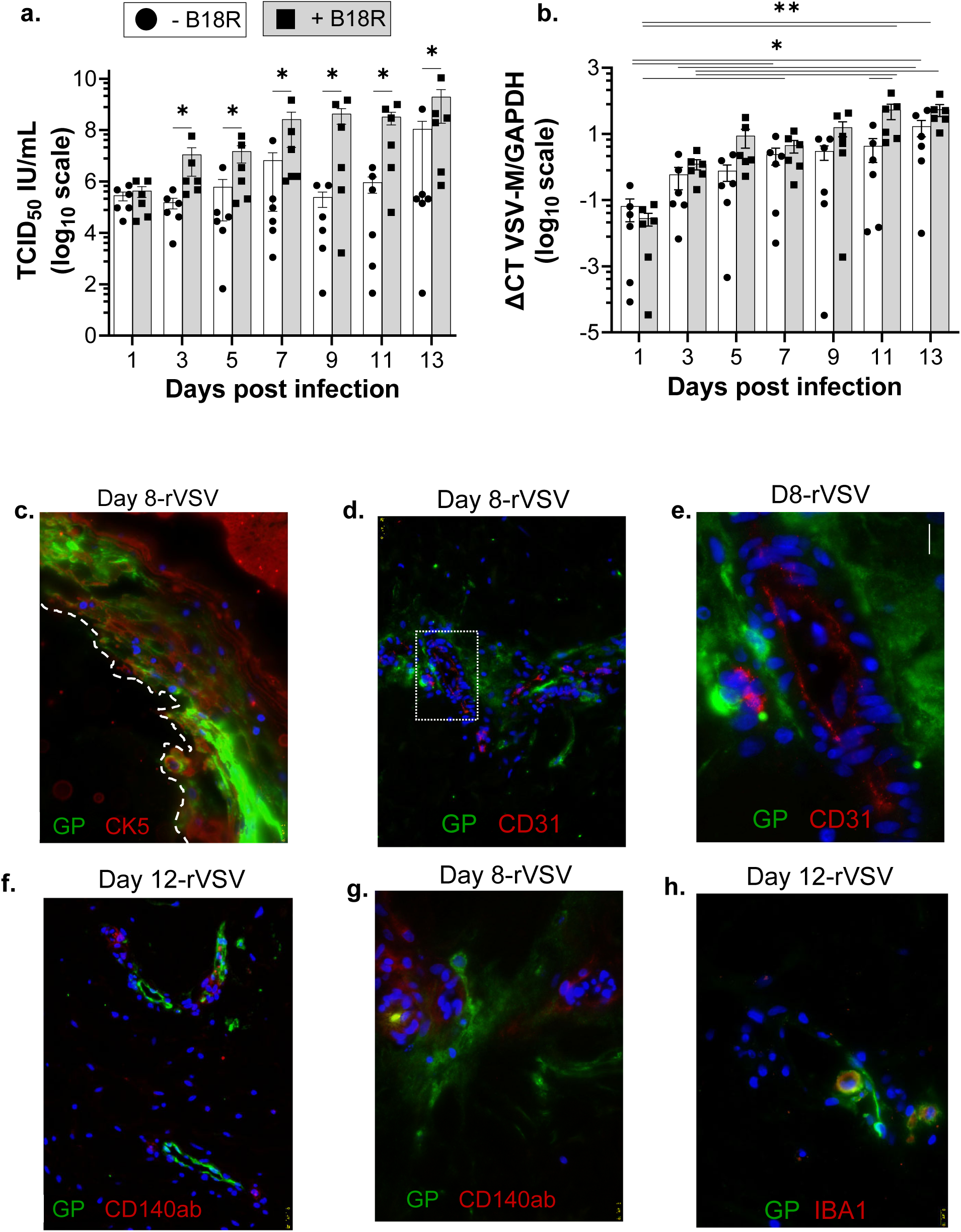
rVSV/EBOV GP tropism is similar to EBOV in human skin explants. Human skin explants were generated and maintained as in Figure 1. Explants were inoculated with 10^7^ iu of rVSV/EBOV GP per explant. **a-b)** After washing twice to remove input virus, explants were maintained in the presence or absence of 0.1 mg/mL B18R. **a**) Viral titers present in basal supernatants were collected on days indicated and assessed on Vero cells. **b**) In parallel studies, explants that were infected with 10^7^ iu of rVSV/EBOV GP were assessed for virus load by qRT-PCR. RNA was extracted from replicate explants on days indicated and viral load was normalized using the GAPDH housekeeping gene. Data is expressed as mean ± SEM. n = 6 replicates per condition from three donors. **c-h**) Replicate explants were collected on the days indicated and FFPE sections were immunostained for virus (EBOV GP in green), and cell lineage markers CK5, CD31, CD140ab, or IBA1 (red) and mounted with DAPI (blue). Viral staining was strongly expressed by the CK5^+^ epidermal keratinocytes **c**) and surrounded CD31^+^ (**Fig. 4.d, e**) or CD140ab^+^ (**Fig. 4f, g**) dermal structures/cell populations. **h**) IBA1^+^ dermal macrophages were also virus positive. Scale bar = 25 uM for panels d, f and 10 μM for the remaining panels.

Consistent with the BSL4 studies, immunostaining identified virus-positive epidermal and dermal cell subsets after rVSV/EBOV GP infection of skin explants (**Fig 4)**). Typically by day 8, strong focal viral staining was observed in CK5^+^ keratinocytes (**Fig. 4c),** which was often accompanied by deterioration of the epidermal structure (**Suppl. Fig. 3c and d**). In the dermis, viral staining of individual elongated cells, CD140ab^+^ cells suggested that fibroblasts support infection (**Fig. 4f, g, Suppl. Movie 2**). Also virus-positive in the dermis were single cells and structures reminiscent of components of the vasculature (**Fig. 4d and e, Suppl. Fig. 3e-h, Suppl. Movie 3**) or clusters of immune cells (**Fig. 4h, Suppl. Fig 3d**). In total, these findings indicate that our lower containment virus rVSV/EBOV GP has a similar tropism to EBOV in human skin.

### Trafficking of virus to the skin’s epithelial surface

Our evidence that the tropism of EBOV and rVSV/EBOV GP is similar allowed us to more extensively use low containment virus to examine various aspects of tropism in skin explants. To understand the contribution of dermal and epidermal cells to rVSV/EBOV GP titers following basal infection, explants were infected with 10^7^ iu of rVSV/EBOV GP, and the input virus was removed the next day. On the day of harvest, explants were treated with dispase to separate the dermal and epidermal layers, and infectious titers present in the two tissue layers were evaluated. Infectious virus was detected in 6 of 15 dermal samples as early as day 2 (**Fig. 5a**). Over time, titers continued to increase, with all but one dermal sample having detectable titers on day 4 through 14, albeit the levels of virus varied widely between explants. Four of 15 epidermal samples also had readily detected infectious virus on day 2. Titers within the epidermis continued to increase until day 8 when titers plateaued. On a per gram basis, levels of infectious virus produced in the epidermis were consistently higher than the dermis even on day 2. This is partially due to the weight normalization of the tissues and differences in the thickness and cellularity of the epidermis (e.g. A 5 mm punch biopsy yields ∼2.97 mg epidermis and ∼55.36 mg dermis when separated). Our data indicate that the virus traffics from the media through the dermis to more apical regions of the explants and that permissive cell populations support virus production in both skin layers.

**Fig. 5.**
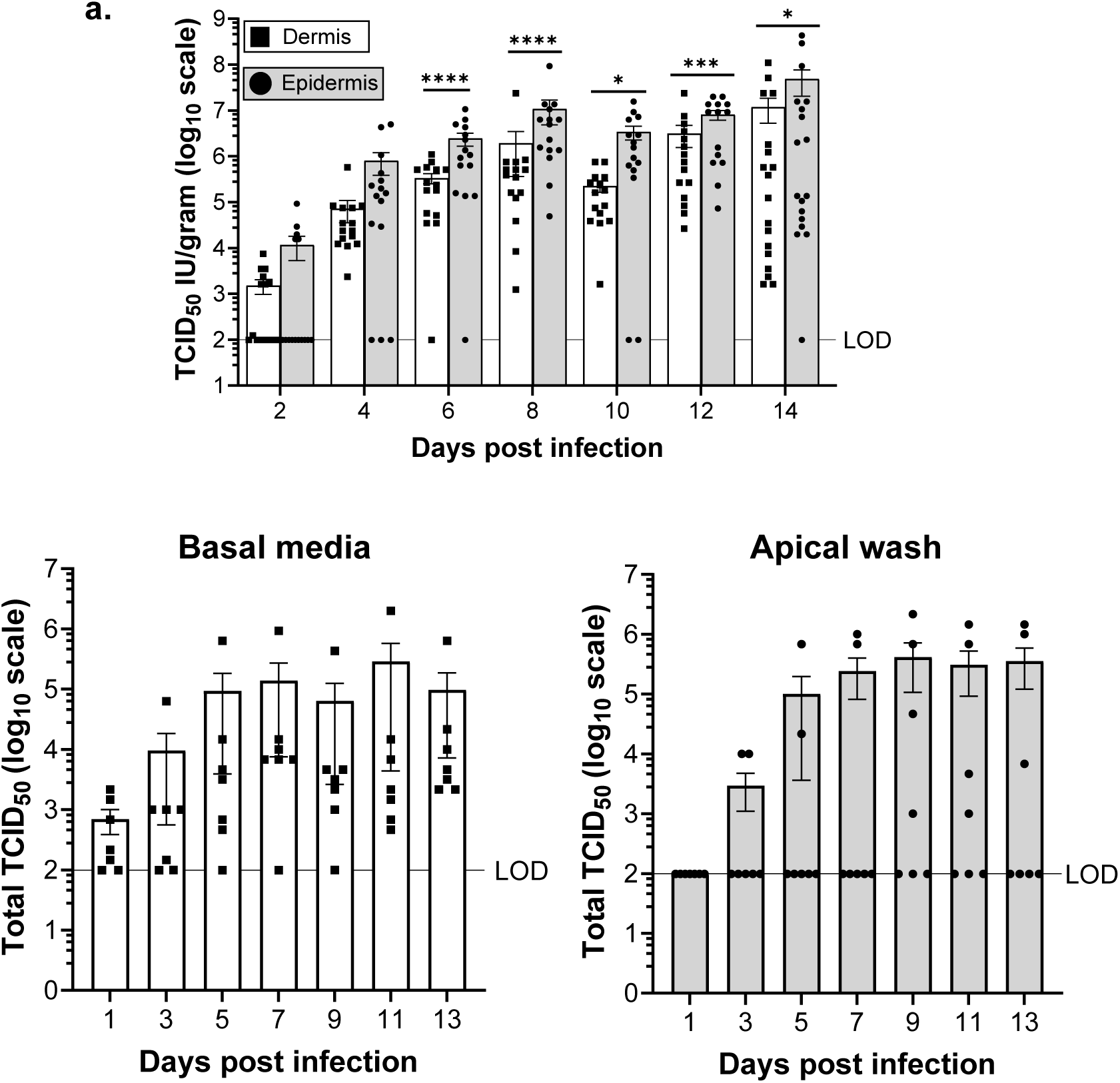
Virus traffics through the explants over time. **a)** Infectious viral titers in the dermis and epidermis over infection. Skin explants were inoculated basally with rVSV/EBOV GP (10^7^ iu) in the presence of ruxolitimib and input virus was removed at 16 hpi. After input virus was removed, explants were washed twice, and then moved to a fresh insert/well with media containing ruxolitimib. On the day of harvest, explants were treated with dispase to separate the dermis and epidermis. Individual tissues were homogenized in PBS, filtered and infectious virus was evaluated on Vero cells (n=4 independent experiments). Data are shown as individual points with the mean ± SEM. Student’s t test, *p<0.05 ***p<0.001 ****p<0.0001. **b**) Infectious virus present in basal supernatants (left panel) or the apical surface of explants (right panel) over the course of infection. Skin explants were infected basally with rVSV/EBOV GP (10^7^ iu) in the presence of ruxolitimib. Input virus was removed, explants were washed twice, and then moved to a fresh insert/well with media containing ruxolitimib. Media was collected and refreshed every other day. At the time indicated, basal supernatants were collected and 10 μL of sterile PBS was placed on the explant stratum corneal surface for one minute and PBS was collected. This was repeated, PBS samples were pooled and titered on Vero cells (n=2 Independent experiments). Data are shown as individual points with the mean ± SEM.

Our findings are the first to demonstrate that keratinocytes within the epidermal layer serve as cellular targets for EBOV infection. We postulated that spread of a virus through the skin layers to the epidermis might lead to the presence of infectious virus on the skin surface, providing a novel route of virus trafficking and transmission. To examine this, we measured EBOV and rVSV/EBOV GP on the apical surface of explants throughout infection. In initial studies, larger (8-10 mm) explants were basally infected with rVSV/EBOV GP for 16 hours. After removal of the input virus, supernatants were collected and the media was completely refreshed every other day. To sample the virus on the apical surface, PBS was pipetted onto the center of the epidermis for ∼ 1 minute. The bubble of PBS was then collected and the sampling was repeated. The PBS was pooled and assayed for infectious virus in comparison to the virus in the basal supernatant (**Fig. 5b**). Consistent with our other findings, the basal media from 5 of the 7 explants contained low but detectable levels of infectious rVSV/EBOV GP at the earliest time point, and these levels increased over time, while the virus was not detected at the apical surface until day 3. At later time points, some of the explants had detectable infectious virus present on the apical surface, with average titers ranging from 10^3^ to greater than 10^5^ in the 20 μL sample. In a single study using EBOV infection, viral RNA could be detected on the epidermal surface by day 7 in some samples (**Suppl. Fig. 4**). The presence of infectious virus on the epidermal surface after inoculation into the basal media indicated that virus can traffic through the explant to the surface of the epithelium.

### Human skin explants are effective multicellular model systems for investigating antiviral therapies

Accessible tissue models for examination and development of new antivirals against filoviruses are limited, with investigations frequently jumping from simple monoculture studies to expensive and multifaceted animal models. Infections of complex tissues are rarely performed outside of the context of animal studies, although with the establishment of organoid cultures (*44*) these kinds of studies are becoming more frequent. As we have shown, human skin explants are highly relevant to EBOV infection, support robust virus infection, and are composed of a complex mixture of different permissive immune and non-immune cell types. Further, these disposed-of tissues are easily obtained from healthy human donors. Hence, these explants may serve as excellent intermediate model systems for characterizing novel antivirals.

To evaluate if these explants serve as robust multicellular intermediate model systems that offer insight into small molecule efficacy in a readily accessible human tissue, we assessed several well-established EBOV inhibitors for their ability to block EBOV or rVSV/EBOV GP infection in skin explants. The cysteine protease inhibitor E64, the endosomal Ca2^+^ channel blocking agent tetrandrine, and the pro-inflammatory cytokine interferon-γ (IFN-γ) were evaluated as they have been shown previously to block infection (*16, 20, 45*). Skin explants were submerged in media containing the appropriate concentration of inhibitor overnight at 37°C to allow tissue penetration. Explants were retrieved and placed in a transwell format and infected via the basal media in the presence of the inhibitor. The input virus was removed the next day. Media containing the inhibitor was refreshed every other day. On day 10 (and other days not shown), rVSV/EBOV GP was readily detected in the supernatant in the absence of inhibitors (**Fig. 6a**), while the inhibitors were effective at blocking rVSV/EBOV GP infection of the explants in a dose-dependent manner, with no significant effect on explant viability (**Suppl. Fig. 5a**).

**Fig. 6.**
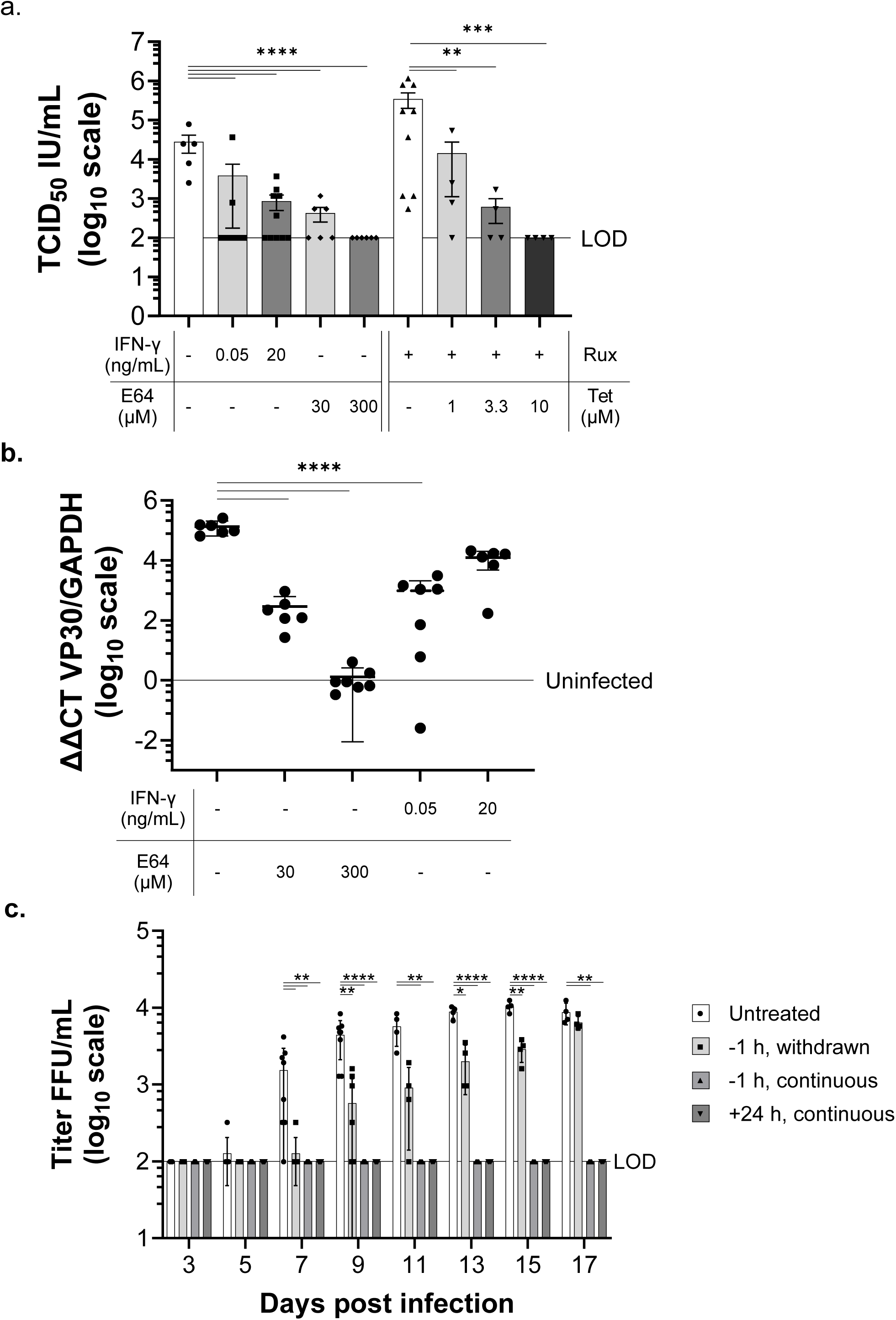
Human skin explants serve as excellent models for drug discovery. **a)** IFN-γ, E64 and tetrandrine (Tet) inhibit production of rVSV/EBOV GP in skin explants. Explants were immersed overnight in the appropriate concentration of drug and then placed in transwells containing the appropriate concentration of inhibitor in the basal media. Skin explants were infected basally with rVSV/EBOV GP (10^7^ iu) in the presence of the inhibitors overnight. Explants were then washed twice and moved to a fresh insert/well with media containing the appropriate concentration of drug throughout infection. Media was collected and refreshed with drug inhibitors every other day. Shown are viral titers detected in basal supernatants on day 10 of infection. Statistical significance was determined by one-way ANOVA on log-transformed values as compared to no drug, infected control. Data is shown as individual points with the mean ± SEM. **b)** IFN-γ and E64 inhibit EBOV replication of skin explants. Studies were performed in a similar manner to panel a, but infected with 10^4^ iu of EBOV-GFP. Shown are viral loads from Day 14 explants that were homogenized in 1 mL of Trizol. RNA was extracted and qRT-PCR performed. Shown are the ΔΔCt values of EBOV VP30 expression normalized for GAPDH expression and divided by the uninfected treatment controls. Uninfected line is defined as the average Ct value of uninfected explants from all treatments divided by itself (uninfected = 1). Statistical significance was determined by one-way ANOVA on log-transformed values as compared to no drug, infected control. Data is shown as individual points with the mean ± SD. **c)** Ability of tetrandrine to delay or abrogate inhibition of EBOV-GFP infection over a 17-day infection. Explants were not immersed in tetrandrine in this experiment, but 10 µM of tetrandrine was added to basal supernatants either 1 hour before or 24 hours after infection initiation. Tetrandrine was either maintained (continuous) or removed (withdrawn) as indicated. Explants were infected with 10^4^ FFU of EBOV-GFP. Statistical significance was determined by two-way ANOVA on log-transformed values as compared to no drug, infected control. Data is shown as individual points with the mean ± SD.

Similar inhibition of EBOV infection of explants was observed with E64 and IFN-γ (**Fig. 6b**). On day 14 of infection, explants were placed in TRIzol, tissues were homogenized and RNA was extracted. EBOV VP30 and host GAPDH expression were assessed by qRT-PCR. The cysteine protease inhibitor, E64, inhibited EBOV infection in a dose-dependent manner. In contrast, the lower dose of IFN-γ significantly reduced VP30 RNA levels, while the higher dose of IFN-γ was less effective. In these studies, GAPDH Ct values were used as a measure of explant viability in the presence of E64 or IFN-γ (**Suppl. Fig. 5b**). In the day 14 explants, GAPDH Ct values of explants treated with the drugs were not distinguishable from explants maintained in media, suggesting that explant viability was not affected by the drugs.

The ability of tetrandrine to block EBOV-GFP was also evaluated. Unlike the other inhibitor studies, explants in these experiments were not immersed overnight in media containing the drug. Instead, media containing 10 μM tetrandrine was present in basolateral media. Tetrandrine was added either one-hour prior to infection or 24 h following initiation of infection. For the one-hour pretreatment, either the drug was maintained for the duration of the experiment or removed before initiation of infection. EBOV-GFP (10^4^ iu) was inoculated into the basolateral supernatant, the input virus was removed one day later, and the media (with or without tetrandrine) was refreshed. Viral titers were assessed in supernatants every other day (**Fig. 6c**). The one-hour drug pre-treatment delayed EBOV production but virus titers recovered by day 17. In contrast, continuous tetrandrine treatment, even if it was initiated 24hpi, was highly effective at inhibiting EBOV infection. In total, these data provide evidence that explants serve as an inexpensive tissue model for testing antivirals against EBOV and we propose these as a useful and highly relevant intermediate step in the exploration of novel antivirals between monocultures and animals.

### EBOV infection of cultured fibroblasts and keratinocytes

Fibroblasts in the dermis and keratinocytes in the epidermis were identified to be virus antigen positive in our skin explants. We postulate that these cells contribute to virus load and egress of virus from an infected individual, yet their interaction with EBOV is not well defined. Hence, we evaluated the ability of cultured primary and immortalized human skin fibroblasts and keratinocytes to support EBOV infection and identified host receptors that mediate virus entry.

Consistent with our explant immunostaining studies, purified primary human skin keratinocytes were permissive for EBOV-GFP and rVSV/EBOV GP infection (**Fig. 7a-c** and **Suppl. Fig. 6a**). Significant viral spread occurred at higher inoculums with infectious virus detectable in the supernatants (**Fig. 7c**). To define host cell receptors present on human keratinocytes that mediate EBOV infection, three primary skin keratinocyte populations were evaluated for gene expression of cell surface receptors known to enhance EBOV binding and internalization into the endosomal compartment. The assessment included a series of C-type lectins such as DC-SIGN and phosphatidylserine receptors such as the TIM family and TAM family. Expression of two of the TAM receptors, Tyro3 and AXL, had consistently higher levels of expression (**Suppl. Fig. 6b**). However, cell surface immunostaining of primary keratinocyte and an immortalized human skin keratinocyte cell line, NHSK-1, demonstrated that Axl was predominantly expressed on the surface of these cells (**Fig. 7d, Suppl. Fig. 6c**). To determine if Axl was required for EBOV infection, infections were performed in the presence of increasing doses of the highly specific Axl signaling inhibitor bemcentinib (*46*). At two different MOIs of rVSV/EBOV GP, we observed decreased virus infection of NHSK-1 cells in the presence of increasing concentrations of bemcentinib at 24 hpi, providing evidence that Axl is critical for virus entry. Finally, to determine if the late endosome/lysosome receptor, NPC1, was also critical for EBOV entry into keratinocytes, we infected NHSK-1 cells that were transfected with NPC1 siRNA which reduced NPC1 present in these cells at 72 hours (**Fig. 7f**). Knockdown of NPC1 expression reduced rVSV/EBOV GP infection (**Fig. 7g**). These studies were performed with or without ruxolitimib. We found that in the absence of ruxolitimib, the irrelevant, scrambled siRNA decreased virus infection, suggesting that transfection of the siRNA was stimulating the innate immune responses of keratinocytes. However, transfection of the NPC1 siRNA reduced virus infection in the absence and presence of the Jak1/2 inhibitor, ruxolitinib. Consistent with an important role of NPC1 in rVSV/EBOV GP entry into keratinocytes, the small molecule inhibitor of EBOV GP binding to NPC1, 3.47(*37*), inhibited virus infection in a dose-dependent manner (**Fig. 7h**).

**Fig. 7.**
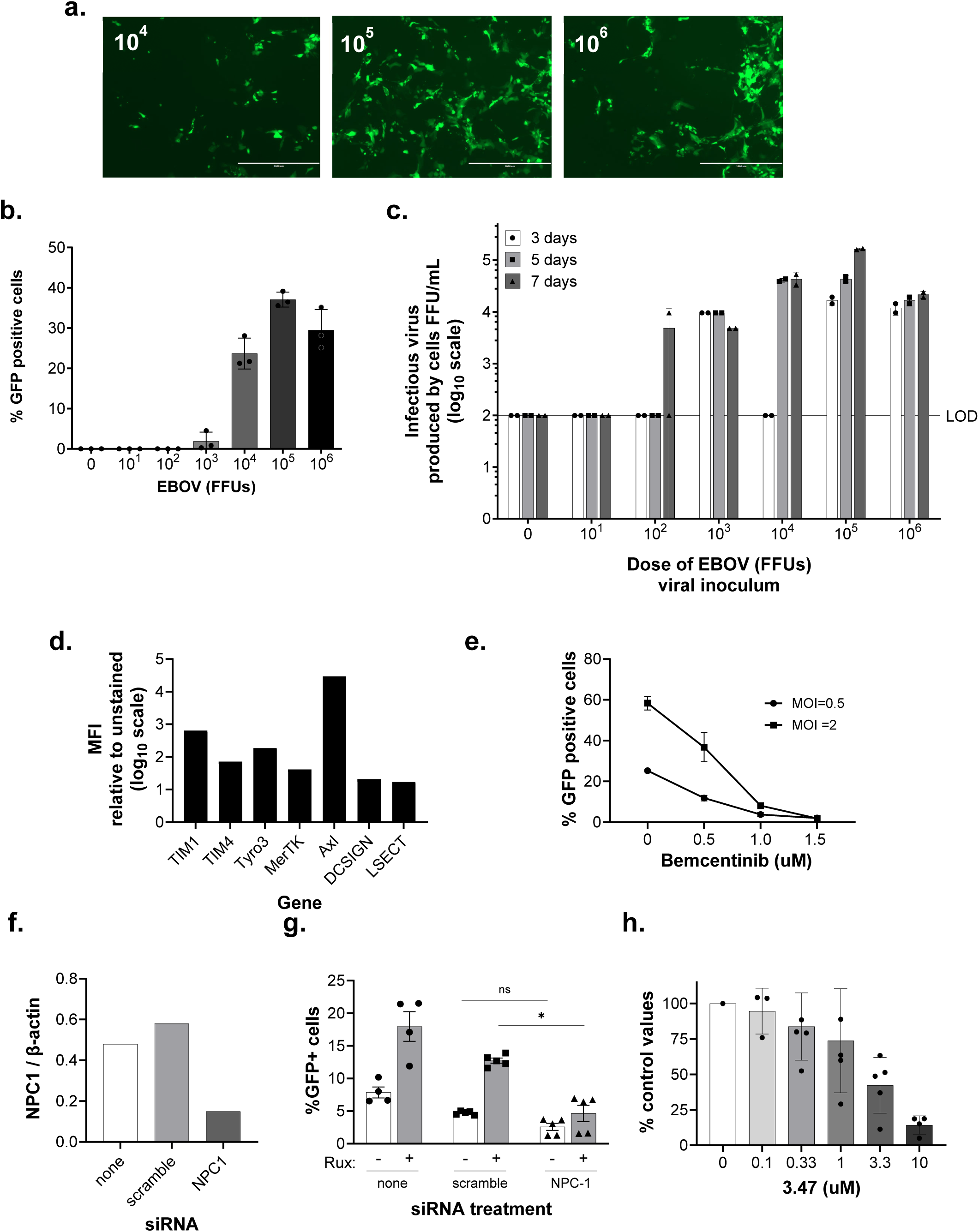
Primary and immortalized human keratinocytes support EBOV infection in an AXL- and NPC1-dependent manner. **a-c**) EBOV-GFP infection of primary skin keratinocytes. Keratinocytes infected with 10^4^-10^6^ iu of EBOV-GFP were imaged on day 5 of infection (a) and quantitated (b). Scale bar = 1 mm. Supernatants were collected every other day on days 3-7 and titered for infectious virus on Vero cells (c). **d-h**) Utilization of receptors by EBOV in the immortalized human skin keratinocyte line, NHSK-1. Cell surface staining of known cell surface receptors utilized by filoviruses (d). Inhibition of rVSV/EBOV GP infection by the Axl signaling inhibitor bemcentinib in a 24 h infection. Data is shown as the mean ± SD of three independent experiments (e). Knock down of NPC1 expression in NHSK-1 cells as evaluated in an immunoblot (f). Loss of NPC1 expression results in decreased rVSV/EBOV GP infection in the presence or absence of rux. Data is shown as individual points with the mean ± SEM (g). Ability of the NPC1 inhibitor 3.47 to block rVSV/EBOV GP infection. Data is shown as the mean ± SD of two or three independent experiments (h).

Similar studies were performed with primary and immortalized human skin fibroblasts (NHSF-2) with surprisingly similar results to that found in keratinocytes since a number of different cell surface receptors can mediate EBOV uptake into cells (*47*). Primary skin human fibroblasts were infected with EBOV and a dose-dependent increase of virus infection was evident on day 7 (**Fig. 8a and b**). At doses of 10^3^ FFU or higher, production of infectious virus in the supernatant of infected cells was evident (**Fig. 8c**). Primary fibroblasts also supported rVSV/EBOV GP infection (**Suppl. Fig. 7**). Gene expression studies in the NHSF-2 cells demonstrated that Axl was more than 100-fold more highly expressed than other known cell surface receptors used by EBOV (**Fig. 8d**) and treatment of NHSF-2 cells or a primary skin fibroblast population with the Axl inhibitor, bemcentinib, inhibited virus infection (**Fig. 8e**), indicating that Axl is critical for EBOV entry into fibroblasts. Finally, the requirement for the endosomal receptor NPC1 was evaluated in siRNA knock down and 3.47 inhibition studies. As anticipated, decreased NPC1 expression or treatment with 3.47 resulted in decreased EBOV GP-dependent entry into NHSF-2 (**Fig. 8f, g and h**). In total, these studies demonstrate that both skin fibroblasts and keratinocytes support EBOV infection in an Axl and NPC1-dependent manner.

**Fig. 8.**
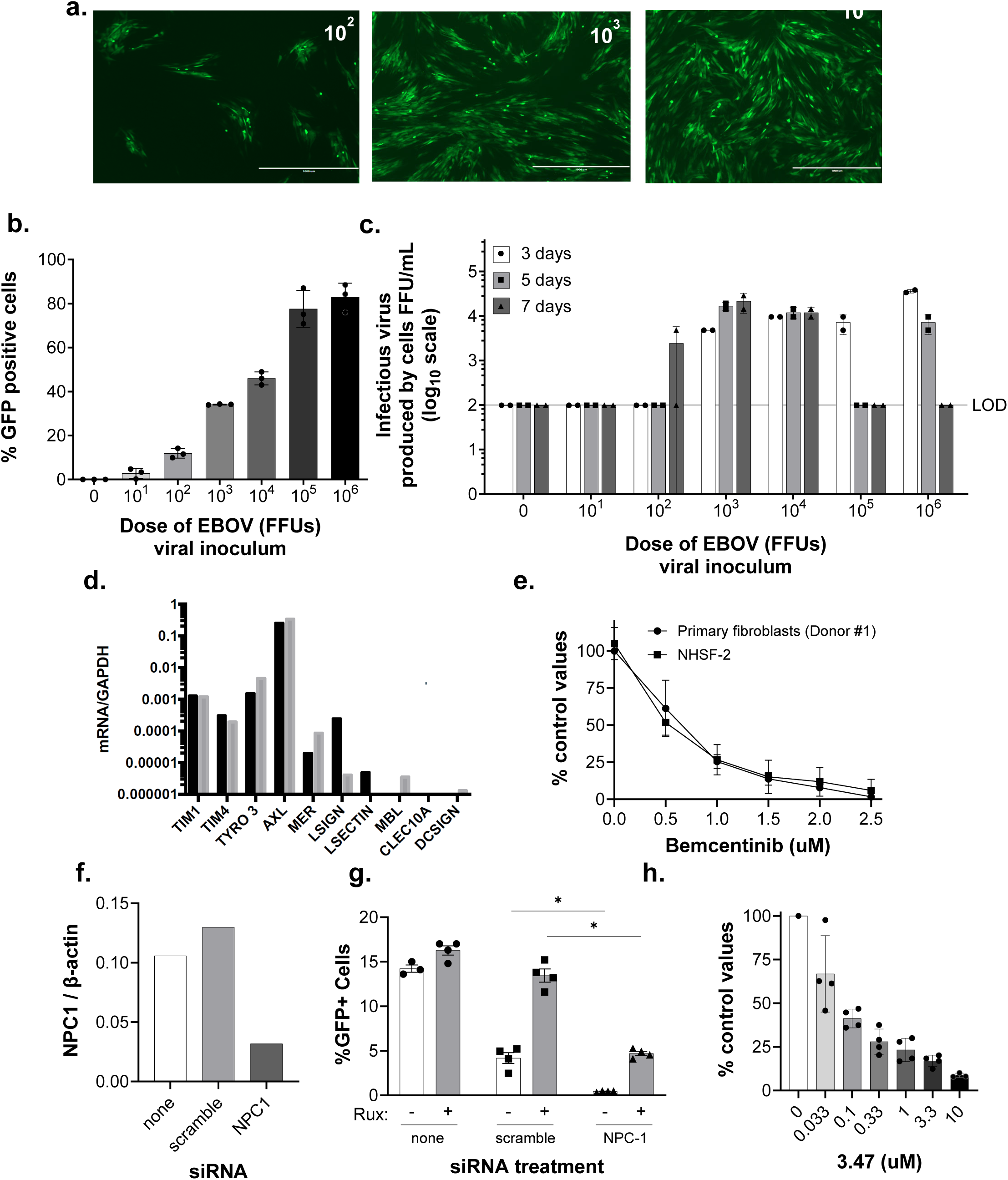
Primary and immortalized human skin fibroblasts support EBOV infection in an AXL- and NPC1-dependent manner. **a-c**) EBOV-GFP infection of primary skin fibroblasts infected with 10^2^**-**10^4^ iu of EBOV-GFP were imaged on day 5 of infection (a) and quantitated (b). Scale bar = 1 mm. Supernatants were collected every other day on days 3-7 and titered for infectious virus on Vero cells (c). **d-h**) Utilization of receptors by EBOV in the immortalized human skin fibroblast line, NHSF-2. Gene expression of characterized cell surface receptors utilized by filoviruses (d). Inhibition of rVSV/EBOV GP infection by the Axl signaling inhibitor bemcentinib in a 24 h infection. Data is shown as the mean ± SD of three independent experiments (e). Knock down of NPC1 expression in NHSF-2 cells as evaluated in an immunoblot (f). Loss of NPC1 expression results in decreased rVSV/EBOV GP infection in the presence or absence of ruxolitimib. Data is shown as individual points with the mean ± SEM (g). Ability of the NPC1 inhibitor 3.47 to block rVSV/EBOV GP infection. Data are shown as the mean ± SD of three independent experiments (h).

## Discussion

The skin serves as a physical barrier to entry and egress of pathogens (*48*). There is anecdotal and experimental evidence indicating that that infectious virus on the skin surface is one route of person-to-person transmission (*7, 8, 12*). While skin petechiae are observed in some filovirus-infected individuals as well as NHP (*11*), not all EBOV infections are associated with skin petechiae (*27, 49*). This suggests that one route of virus trafficking to the skin’s surface may be via infection of skin cells rather than direct transmission of virus from the blood. Prior immunohistochemistry studies identify virus antigens present in skin tissue (*6, 7, 11, 12, 49, 50*), yet previous co-staining of cell markers with EBOV antigen had not been performed. Therefore, cells within the skin responsible for transmission of the virus had not been definitively identified.

Here, we used human skin explants to demonstrate that dermal and epidermal cells support EBOV infection. Use of the human skin explant model has several advantages in EBOV studies, including that: 1) de-identified human skin is readily available from surgeries, 2) it is a complex mix of cells from an organ that is relevant at late times of EBOV infection and 3) it readily allows investigation of EBOV infection in tissues from a broad cross-section of the population, providing insights into host genetic heterogeneity. Skin explants supported robust levels of virus infection over more than two weeks with little to no evidence of reduced cell viability in the uninfected or infected explant cultures. Administration of the virus to the basal surface of the dermis in a transwell format resulted in a detectable virus in the dermal and epidermal tissue and on the apical surface of the epidermis, indicating movement of the virus through the different skin layers. Surprisingly, on a per-gram basis, virus replication was more robust in the epidermal layer than within the dermis. The virus rapidly trafficked through the dermis to the epidermis and production of infectious rVSV/EBOV GP could be detected in the epidermis as early as day 3. We postulate that virus infection of the epidermis results in the trafficking of infectious viruses to the skin’s surface. Our studies provide a new paradigm for understanding filovirus egress from the body and a mechanism of transmission from one host to the next.

Several different cell types were found to support EBOV infection in the skin explants. Within the dermis, we observe IBA1^+^ macrophages, CD31^+^ endothelial cells, and CD140ab^+^ fibroblasts expressing virus antigens. Tissue macrophages are well established to support EBOV infection in a variety of tissues (*28, 29, 51–53*). While fibroblasts and other related stromal cells that form a tissue matrix are postulated to support EBOV infection(*27*), to our knowledge, this is the first demonstration of co-staining of EBOV with a fibroblast marker. We also report for the first time that CK5^+^ keratinocytes support infection. The production of infectious virus in these cells likely contributes to the infectious virus present on the surface of explants.

These explants served as excellent tissue model for examining the profiles of potential antivirals. We show that the small molecule inhibitors, tetrandrine, and E64, as well as the proinflammatory cytokine, IFN-γ, blocked EBOV and rVSV/EBOV GP infection in the explants. IFN-γ receptors are present on fibroblasts in the dermis and keratinocytes in the epidermis as well as myeloid populations, consistent with an important role of these cells in skin infection (*54, 55*). We propose that this easily accessible and relevant tissue should be incorporated into antiviral workflows as a highly useful step before animal studies to evaluate drug efficacy.

Of note, we do see variation in the robustness of infection between explants generated from different donors. While explants from almost all donors became infected, the production of infectious virus ranged from ∼10^3^ to 10^6^ and was donor-dependent. This is not surprising given the outbred nature of humans.

Purified human skin fibroblasts and keratinocytes were found to support EBOV infection, consistent with our findings that these cell types are viral antigen-positive in our infected explants. Both cell types required the cell surface receptor Axl and the endosomal receptor NPC1 for EBOV entry. Our prior studies demonstrated that Axl, is abundantly expressed on the surface of endothelial cells and mediates binding and internalization of rVSV/EBOV GP (*56*). We propose that Axl is used to enhance virion binding to these cells and internalize the virus into the endosomal compartment where NPC1 interactions are required. While Axl gene expression was more that 100-fold higher in fibroblasts than expression of the other surface receptors examined, gene expression of Axl and Tyro3 were roughly equivalent in keratinocytes. Surprisingly, given the keratinocyte finding, Axl was the only receptor detected on the surface of keratinocytes. For both cell types, virus infection was effectively inhibited by bemcentinib, a signaling inhibitor that is highly specific for Axl (*57*). Future studies with Axl knock-out mice should provide additional insights into the critical nature of this receptor in supporting filovirus infection of the skin.

EBOV encodes two antagonists of innate immune responses, VP35 and VP24 (*58–62*). The role of these virally encoded proteins on the ability of EBOV to replicate in the skin is an important unanswered question. The innate responses of immune and non-immune cells within the skin to EBOV infection require more examination. Our observations that type I IFN pathway inhibitors, B18R and ruxolitinib, enhance rVSV/EBOV GP infection is consistent with a significant IFN response in our skin explants. Dermal and epidermal cells may elicit antiviral innate immune responses that may help control EBOV transmission through skin contact.

Studies are ongoing to address this important question.

## Materials and Methods

### Procurement of human skin and generation of skin explants

De-identified skin from healthy individuals was obtained within 1-2 hours of surgery at the University of Iowa Healthcare through the Tissue Procurement Core (https://medicine.uiowa.edu/tissueprocurement/about). Most skin samples were taken from breast reduction surgeries, but skin samples were also obtained from panniculectomies and circumcisions. Skin samples were passed through 70% ethanol and povidone-iodine solutions and trimmed of excess adipose tissue. Replicate punch biopsies (3-10 mm) were generated and placed in complete media composed of MEM or high glucose DMEM (ThermoFisher, Waltham, MA) containing 10% fetal calf serum, 1.6 µg/mL gentamycin solution, 200 units/mL of penicillin, 200 μg/mL of streptomycin and 0.5 µg/mL amphotericin b (skin media). For BSL4 studies, skin biopsies were shipped in complete media overnight on wet ice to the Texas Biomedical Research Institute, San Antonio, TX.

Explants were placed dermal side down on the membrane surface of a transwell insert (Corning #3460, 0.4 µm pore size). The inserts were placed within a tissue culture well with sufficient media to maintain the skin explant at the air/liquid interface. Plates were incubated at 37°C, 5% CO_2_ for the duration of the studies.

### Generation of virus stocks and evaluation of viral titers

Vero cells (ATCC CCL-81) or Vero E6 (ATCC CRL-1586) were used to generate EBOV and rVSV/EBOV GP stocks. These cells were passaged in DMEM with 5% or 10% fetal calf serum (FCS), 100 units/mL of penicillin, and 100 μg/mL of streptomycin (1% PS).

All work with replication-competent EBOV was performed in the biosafety level 4 (BSL-4) laboratory at the Texas Biomedical Research Institute, San Antonio, TX, according to standard operating procedures and protocols approved by the Institute’s Biohazard & Safety and Recombinant DNA committees. The recombinant EBOV variant Mayinga expressing GFP (EBOV-GFP, NCBI accession number KF_990213) was grown in Vero cells for 7 days. The culture supernatant containing the virus was clarified of cell debris by centrifugation at 3,500 rpm for 20 min, then overlaid above a 20% sucrose cushion in PBS, and ultracentrifuged at 4°C for 2 hours at 28,000 rpm. The virus pellet was resuspended in PBS and the virus titer was determined by incubating serial dilutions of the stock on Vero cells for 24 hours. Cells were incubated with Hoechst dye (Thermo Fisher Scientific) to stain nuclei, then photographed by a Nikon system, and analyzed by CellProfiler (Broad Institute, Cambridge, MA) to quantify nuclei and infected (GFP-positive) cells.

To generate the rVSV/EBOV GP stocks, Vero or Vero E6 cells were infected with a low MOI (∼0.005) of infectious, recombinant vesicular stomatitis virus (rVSV) encoding EBOV GP and GFP in place of the native G glycoprotein. Supernatants were collected after ∼24 to 36 hours when notable cytopathic effect was evident. Supernatants were centrifuged at 1500 x g to remove cell debris and filtered through a 0.45 μm filter. Virus in the supernatants was layered over a 25% sucrose/PBS cushion and ultracentrifuged at 120,000 x g. The resulting pellets were thoroughly resuspended in PBS, aliquoted, and frozen at −80°C until use. To determine the tissue culture infectious dose (TCID_50_) of the stock, 12,000 cells were plated in a 96-well format and the virus was serially diluted on the plate (n=8 wells/dilution). Infections were maintained for 5 days, and GFP-positive (GFP^+^) cells were assessed. The Reed and Muench method was used to determine the TCID_50_ of the stock (*15*). In some endpoint assays, titers of infectious rVSV/EBOV GP were quantitated by serial dilutions of virus-containing samples on 12,000 Vero or Vero E6 cells for 24 hours at 37°C. Cells were lifted with trypsin (%) and GFP^+^ cells were quantitated by flow cytometry.

### Viral load determinations

EBOV load in supernatants, PBS, or explants was determined by qRT-PCR. Briefly, infected samples were placed in 1 mL of Trizol LS (ThermoFisher, #10296010) and stored at - 80° C until use. rVSV/EBOV GP samples were placed in 0.5 mL of Trizol (ThermoFisher, #15596026). RNA was isolated per the manufacturer’s instructions. Total RNA was quantified and 1 μg was converted to cDNA using the High-Capacity cDNA Reverse Transcription kit (ThermoFisher, #4368814). qPCR was performed using PowerUp™ SYBR™ Green Master Mix (Applied Biosystems/Thermo Fisher Scientific, #A25742) as directed. Twenty ng of cDNA was amplified in duplicate per sample using a QuantStudio™ 3 Real-time PCR machine (Applied Biosystems (ABI)). Threshold cycle (Ct) values were averaged and normalized to the housekeeping gene, GAPDH, to calculate the ΔCt value. Alternatively, Ct values in supernatants and PBS samples were compared to a standard curve. A standard curve consisting of serial dilutions of the VP30 plasmid was used to determine copy number: copy number = (concentration of plasmid (ng) x 6.022×10^23^)/((1451*660g/mol) x (10^9^ ng/g)). qPCR was performed on the VP30 plasmid dilutions to determine the average Ct value. The copy number and Ct values were used to generate a logarithmic standard curve and the equation from the standard curve was applied to samples where specified. The primers utilized in these studies are listed in **Table 1**.

**Table. 1.**
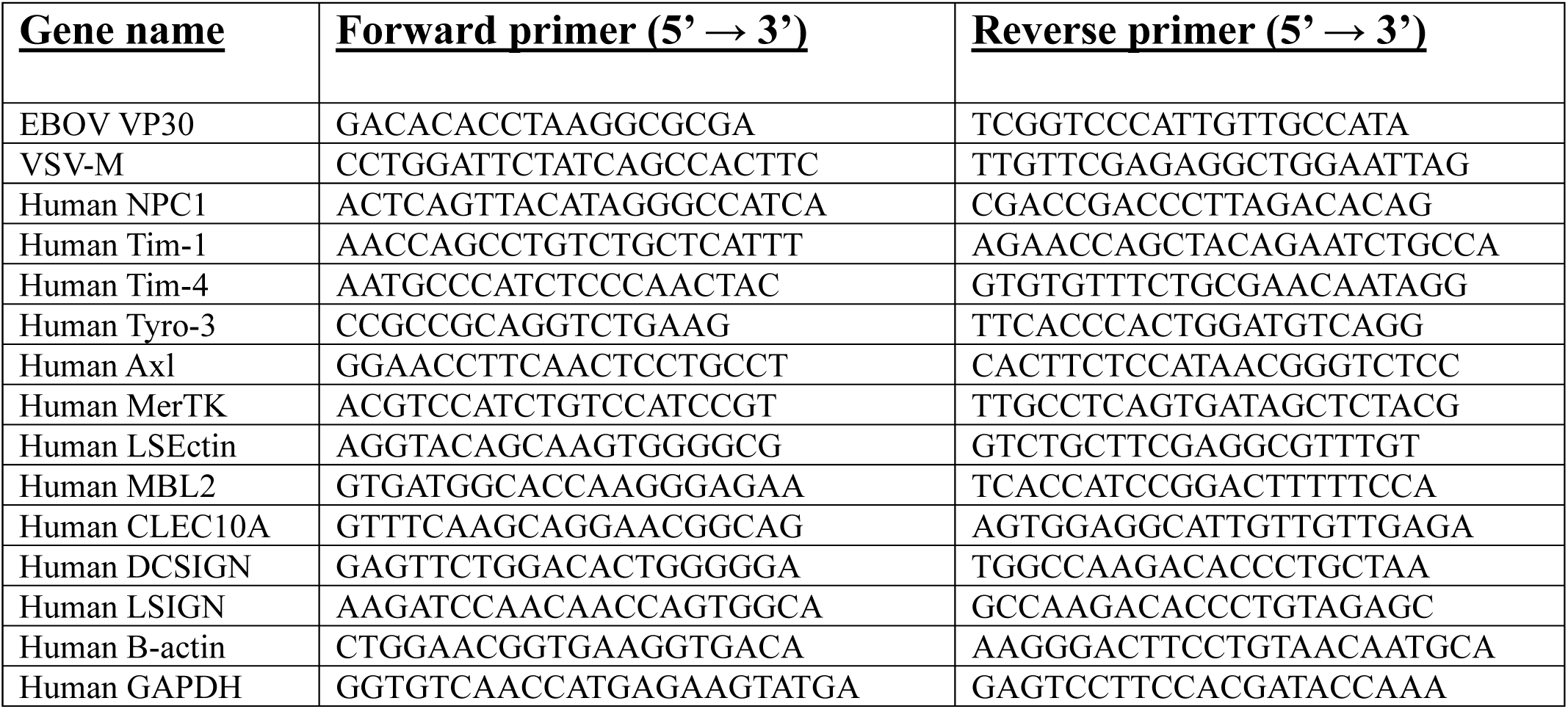
Oligonucleotides primers used in this study.

### Skin explant infections

Virus was added to the basal media within 24 hours of culture initiation and replicate “mock infection” explants were cultured in parallel without viral inoculation. For each experiment, the dose of virus, the timing of exposure, culture duration, and the samples collected/assays run are described in the figure legend. Some rVSV/EBOV GP infections were performed in the presence of 200 ng/mL of the type I interferon (IFN) binding protein, B18R (Invivogen, #inh-b18r), to block endogenous IFN activity or 1 μM of the Jak1/2 inhibitor ruxolitinib (rux) (Selleckchem, #S1378) to block IFN signaling. In some cases, the input virus media was removed after 7-16 hours by two 1X PBS washes of the well before media replacement. Alternatively, 7-16 hours after the input virus was added, the explants were transferred to new plates and washed two times with 1X PBS and then transferred to new wells containing fresh media to eliminate residual input virus. Supernatants were collected on days indicated, filtered through a 0.45 μm filter, and stored at −80°C.

Explants collected at various days post-inoculation were: 1) fixed in formalin before paraffin embedding for immunostaining studies, 2) homogenized in PBS for titering, or 3) placed in TRIzol (ThermoFisher, #15596026) for viral load assessments. In some studies, the explant dermis and epidermis were separated by digestion in dispase II (4 mg/mL) for 2 h at 37°C. The separated tissues were mechanically homogenized with a microcentrifuge pestle in 500 µL of 1X DPBS and stored at −80°C until titered. EBOV-infected biopsies were fixed in formalin for one week before removal from the BSL-4 facility per approved protocols.

### Impact of antivirals on virus infection of skin explants

EBOV infection is inhibited by the cysteine protease inhibitor, E64, the Ca2^+^ channel blocker, tetrandrine, and interferon-gamma (IFN-γ)(*16–20*). To test their effect in our skin explant model, antiviral agents were diluted in KSFM or skin media and used at concentrations noted in figure legends. Explants were pre-incubated (submerged) in the noted concentration of inhibitor diluted in media overnight at 37°C. Explants were placed on fresh transwell inserts and maintained with the appropriate inhibitor in the media below. For EBOV-GFP infection, each explant was infected with 10^4^ iu for 16-18 hours. For rVSV/EBOV GP studies, each explant was infected with the concentration of virus noted in the figure legend (usually 10^7^ iu) for 16 hours. Explants were washed 2x (1X DPBS containing 2% P/S, 1.6 µg/mL gentamycin, and 0.5 µg/mL amphotericin B) to remove input virus and transferred to a new transwell insert with skin media containing the noted concentration of each antiviral agent in the well below. Supernatants were collected every other day for viral quantitation as above and LDH levels were assessed (below).

### Immunofluorescent staining and microscopy of skin explants

Sections (6 µM) of formalin-fixed paraffin-embedded (FFPE) skin explants mounted on glass slides were warmed in a 50°C oven for 45 minutes, deparaffinized with xylene, and rehydrated through incubation in decreasing concentrations of ethanol diluted in H_2_O. Epitope retrieval consisted of 10 mM citrate buffer and 3 cycles (3.5 minutes) of warming to 50-60°C. After cooling to room temperature, tissue was permeabilized with 0.5% Triton X-100 diluted in TBS staining buffer (TBS/Ca/0.09% NaN_3_). Sources, optimal concentrations, and conditions used for staining with each antibody are shown in **Table 2**. Slides were washed and blocked (5% BSA in staining buffer), sections were demarcated with a barrier pen, and primary antibodies diluted in blocking buffer (50μL) were added to each section and incubated in a humidified chamber. In some cases, serial incubation with primary antibodies was necessary to achieve optimal staining. Slides were washed 2x and sections were incubated (37°C for 45 minutes) with species-specific secondary antibodies diluted in blocking buffer (50 μL) (**Table 2**) Slides were washed 2x, dipped briefly in H_2_O, and coverslips were mounted with media containing DAPI (ThermoFisher, # P36934). Images and movies were collected using Leica SPE confocal microscope and Imaris software at the Neural Circuits and Behavior Core. In some cases, color adjustments were made to an entire image, or images were cropped using Adobe Photoshop software.

**Table 2.** Antibodies used in immunostaining studies.

| **<u>Antibody</u>** | **<u>Company, Catalog #</u>** | **<u>Host species</u>** | **<u>Dilution and conditions used</u>** | **<u>Incubation temperature</u>** |
| --- | --- | --- | --- | --- |
| *Primaries:* |  |  |  |  |
| EBOVGP | IBT Bioservices, Cat#0201-020 | mouse | 1:100, overnight | 4°C |
| VP40 | IBT Bioservices, Cat#0201-016 | mouse | 1:100, overnight | 4°C |
| Cytokeratin 5 | ThermoFisher Cat#PA5-29670 | rabbit | 1:200, 2 hours | RT |
| CD31 | ThermoFisher Cat#PA5-16301 | rabbit | 1:100, overnight | 4°C |
| IBA1 | FIJIFILM WAKO, Cat# 019-19741 | rabbit | 1:300, 2 hours | RT |
| CD140ab (PDGFRab) | Abcam Cat#ab32570 | rabbit | 1:100, 2 hours | RT |
| MHC class ii | ThermoFisher Cat#14-5321-82 | rat | 1:100, overnight | 4°C |
| MCM2 | Abcam Cat#ab108295 | rabbit | 1:100, 2 hours | RT |
| *Secondaries:* |  |  |  |  |
| Goat anti-mouse AF488 | ThermoFisher Cat# A11029 | goat | 1:800, 1 hour | RT |
| Goat anti-rabbit AF555 | ThermoFisher Cat# A32732 | goat | 1:800, 1 hour | RT |
| Goat Anti-rat AF555 | ThermoFisher Cat# A21434 | goat | 1:800, 1 hour | RT |
| RT = room temperature |  |  |  |  |

### Explant viability determinations

Explant viability over 14 days was determined through evaluation of lactate dehydrogenase (LDH) in the culture media as directed, except the stop solution was not used (Promega, Madison, WI, CytoTox-One). Briefly, 25 µL of culture supernatant were added to 475 µL of LDH storage buffer (200 mM Tris-HCl, pH 7.3, 10% Glycerol, 1% BSA) and stored at - 80°C until use. Assay controls included an LDH background, a media control, and a maximal LDH release, made by submerging an explant in a 0.04% Triton X-100 solution. A standard curve was performed for each assay using purified LDH from rabbit muscle according to the manufacturer’s instructions (Promega, Madison, WI, LDH-Glo Assay, # J2380). 50 µL of samples, controls, and the standard curve were plated in triplicate on a white opaque, flat bottom 96-well plate (Corning, Corning, NY, REF#353296). 50 µL of CytoTox-ONE reagent was then added to each well and plates were shaken for 10 minutes in the dark at room temperature (RT). After incubation, the fluorescence was read with excitation at 560 nm and emission at 590 nm.

### Primary skin fibroblast and keratinocyte populations and immortalized lines

#### Isolation and characterization of primary skin fibroblasts and keratinocytes

Primary fibroblasts and keratinocyte cultures were obtained from skin tissue of young, healthy adults undergoing surgery for macromastia using methods similar to those described(*21, 22*). Briefly, the skin was disinfected as above and the dermis and epidermis were separated by overnight dispase II treatment (Roche, #4942078001, 10 mg/mL in HBSS, 4°C). Fibroblasts were obtained from minced dermal tissue by overnight collagenase digestion (Worthington, #LS004196, 1000 units/mL in DMEM, 37°C). After complete digestion, the solution was diluted 2-fold in warm 10% FBS/DMEM with antibiotics and centrifuged at 200 x g for 10 minutes. Using the same media, the pellet was washed and the cells were plated and passaged 1:4 with media change every other day. Infections were carried out between passages 3 and 6.

Keratinocytes were isolated from the minced epidermis that was further digested with 0.05% trypsin/EDTA for 10 minutes at 37°C followed by trituration to remove basal keratinocytes. Cells were washed with 2% FBS in PBS and centrifuged at 200 x g for 10 minutes and cultured in keratinocyte serum-free media (KSFM, ThermoFisher, #17005042), which inhibits the growth of non-keratinocyte cells, resulting in a pure population of keratinocytes after passaging. The media was replaced every other day and cells passed 1:4 every 4 or 5 days. Cells were expanded and frozen down at early passage. Infections were carried out between passages 3 and 6.

#### Production and characterization of immortalized skin fibroblasts and keratinocytes

TERT-immortalized human skin fibroblasts (TERT-N-HSF-2) and TERT-immortalized human skin keratinocytes (TERT-N-HSK-1) were generated by retroviral transduction of the reverse transcriptase component of telomere, TERT, as previously described (*21, 22*). Fibroblasts were maintained in 10% FBS/DMEM and keratinocytes were maintained in KSFM and media was changed every other day.

RNA expression of well-established surface receptors for filoviruses was evaluated in primary human skin fibroblasts and keratinocytes. Reverse transcription and qPCR were performed as described above for the detection of viral load. Primers used in these studies are listed in **Table 1**. For primary keratinocytes and our immortalized keratinocyte line, we also assessed cell surface expression of many of these receptors. Assessment of surface receptor expression on primary or immortalized skin keratinocytes was performed by incubating ∼200,000 cells with 1 µg/mL of antibody specific for TIM1 (R&D, #AF1750), TIM4 (R&D, #AF2929), Tyro3 (R&D, #AF859), MerTK (R&D, #AF891), Axl (R&D, #AF154), DC-SIGN (R&D, #MAB1621) and L-SECTin (R&D, #AF2947), or control antibody (R&D, #AB108C) for 30 minutes at 4°C. Cells were washed with 1X PBS x5, then incubated with 1:1000 dilution of Fluorescein (FITC)-conjugated anti-goat or anti-mouse IgG (Jackson Immuno, #705095147 or #715096150) for 30 minutes at 4°C. Cells were washed five times with 1X PBS, then evaluated for FITC intensity on a Beckman Cytoflex instrument.

### Infection of cells in culture

#### EBOV-GFP

Primary keratinocytes or fibroblasts cultured in 96-well plates at ∼30,000 cells/well were incubated with 10^4^, 10^5^, or 10^6^ FFU (keratinocytes) or 10^2^, 10^3^, or 10^4^ FFU (fibroblasts) of EBOV-GFP in DMEM containing 1% FBS, in triplicate, for 16-18 hours. Subsequently, virus inoculum was removed, and cells were washed and overlaid with fresh media for 2, 4, or 6 days. Supernatants were collected and titrated onto Vero cells for 24 hours. All samples were stained with Hoechst dye and photographed and analyzed as described above.

#### rVSV/EBOV GP

All primary or immortalized fibroblasts and keratinocytes were cultured in duplicate or triplicate in a 48-well plate at ∼60,000 cells/well in a volume of 250 - 300 μL of appropriate media. Infections were performed as described in figure legends. At the indicated times, cells were detached with 0.05% Trypsin-EDTA (ThermoFisher, #25300054), washed in 1XPBS, 5% newborn calf serum, and resuspended in 1XPBS with 2% calf serum with azide (FACS buffer). Flow cytometric analysis was done on Becton Dickenson FACSVerse, BD Calibur, or Beckman Cytoflex to quantify the frequency of GFP^+^ cells. At least 3 independent experiments were conducted.

### Inhibitor studies and siRNA knockdown studies in skin fibroblasts and keratinocytes

#### Bemcentinib inhibition studies

TERT-N-HSK-1 cells, primary skin fibroblasts or TERT-N-HSF-2 cells were plated at 50,000 cells/well in KSFM or DMEM with 10% FCS as noted above. Appropriate concentrations of the Axl signaling inhibitor, bemcentinib (kind gift of Dr. James Lorens, University of Bergen) were added to culture media. Keratinocyte cultures were infected with rVSV/EBOV GP at a MOI = 0.5 or 2 (as titered on Vero cells), whereas fibroblast infections were performed at a MOI of 4.6. At 24 hours post infection, cells were lifted with 100 μL of trypsin. Once cells were no longer adherent, 100 μL of new-born calf serum was added to each well and GFP^+^ cells were evaluated by flow cytometry.

#### 3.47 inhibition studies

The NPC1 inhibitor, 3.47, is an adamantane dipeptide piperazine derivative and a kind gift from Dr. James Cunningham, Harvard University. TERT-N-HSK-1 cells or TERT-N-HSF-2 cells were plated at 50,000 cells/well in KSFM or DMEM with 10% FCS, respectively.

Appropriate concentrations of 3.47 was added to cell cultures prior to infection with rVSV/EBOV GP at an MOI of 4.6. At 24 hours post infection, cells were lifted with 100 μL of trypsin. Once cells were no longer adherent, 100 μL of new-born calf serum was added to each well to inhibit trypsin activity and GFP^+^ cells were evaluated by flow cytometry.

#### siRNA knockdown studies

TERT-N-HSF-2 (fibroblasts) or TERT-N-HSK-1 (keratinocytes) were plated at a density of ∼500,000 cells per well of a 6-well plate. Cells were transfected 24 h later with NPC1-specific siRNA or a non-targeting siRNA at a concentration of 25 pmol per well using the Lipofectamine RNAiMAX transfection reagent (Invitrogen, Carlsbad CA). NPC-1 DsiRNA: 5’-GACUUUAUUGACGCUCUG – 3’, 5’-CUUUCUUCAGAGCGUCAA −3’ (IDT, Coralville IA). Non-targeting DsiRNA from IDT was utilized as negative control (IDT, #51011404). Briefly, Lipofectamine RNAiMAX and siRNA were mixed according to the RNAiMAX protocol in serum free media (Opti-MEM for TERT-N-HSF-1, K-SFM for TERT-N-HSK-2) and incubated for 5 minutes at room temperature. Media was removed from cells, washed once with indicated serum free media, then 300 µL of transfectant was added directly to the cells. After a 6 h incubation at 37°C at 5% CO_2_, transfectant was removed from cells and replaced with complete media. 48 h post-transfection, cells were split into a 48-well format at a density of ∼60,000 cells per well in triplicate for infection and remaining cells placed back into 6 well format to validate knockdown via immunoblot. At 72 h post-transfection, cells in 48 well format were infected with rVSV/EBOV GP at an MOI= 10 (keratinocytes) or MOI=1 (fibroblasts) and assessed for GFP^+^ cells 24 h later as described above. In parallel, 72 h post-transfection cells in 6 well format were harvested in 1% SDS to assess knockdown efficiency via immunoblot (Abcam, #ab134113).

### Statistical analysis

Statistical analysis was completed in GraphPad Prism v9.0.2 (GraphPad Software, San Diego, CA). Graphs on a log scale had statistics (including Student’s T test, one-way ANOVA, and two-way ANOVA analyses) performed on log transformed data. Flow cytometric analysis was done in FlowJo v.10.7.1 (Becton, Dickinson & Company, Ashland, OR). Statistical significance was defined as p < 0.05 and denoted by a single asterisk (*) where indicated. More specific details regarding statistical tests and exact n/group can be found in the corresponding figure legends. All presented data is representative of n = 3 independent experiments or as noted in the figure legend.

## Acknowledgements.

The following grants are acknowledged for their support of the project:

National Institutes of Health grant R01AI134733 (WM)

National Institutes of Health grant R21AI144215 (WM)

National Institutes of Health grant UH2AI169710 (WM)

National Institutes of Health grant R21AI154336 (OS)

National Institutes of Health grant R21AI151717 (OS)

National Institutes of Health grant UC7 AI095321 (RAD).

AF was supported by Department of Dermatology, University of Iowa.

## Author contributions

Conceptualization: WJM, KNM, AK, RAD

Methodology: PTR, AK, KNM, WJM. OS

Investigation: PTR, AF, MD, RAP, SC, JE, TPC, PMG, FG and JAD

Visualization: KNM, PTR, JE, WJM

Supervision: WJM, KNM, RAD, AK

Writing—original draft: WJM

Writing—review & editing: WJM, KNM, PTR, RAD, OS

## Data availability

All data needed to evaluate the conclusions in the paper are present in the paper and/or the Supplementary Materials.

---

**Suppl. Fig. 1.**
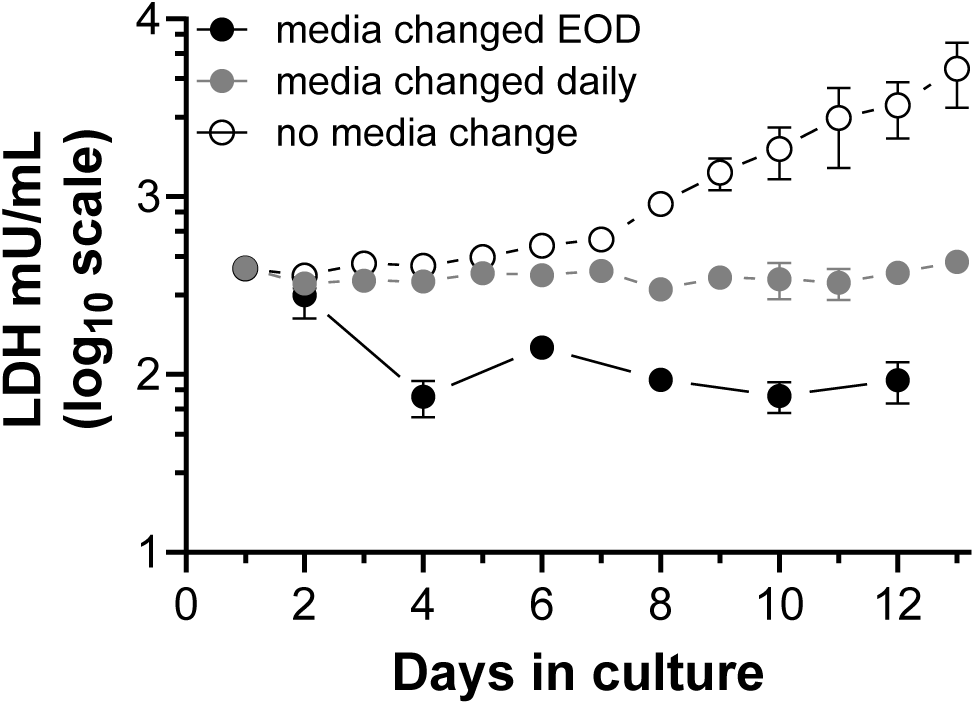
Human skin explant model of EBOV infection: LDH levels in supernatant of uninfected human skin explants under different culture conditions. As an index of viability, lactate dehydrogenase activity (LDH) was measured in supernatants of uninfected skin explants cultured for 13 days without any media change (open circles), with complete media change every day (grey circles), or every other day (black circles). Data are expressed as mean ± SD. n = 6 replicates per condition from a single donor.

**Suppl. Fig. 2:**
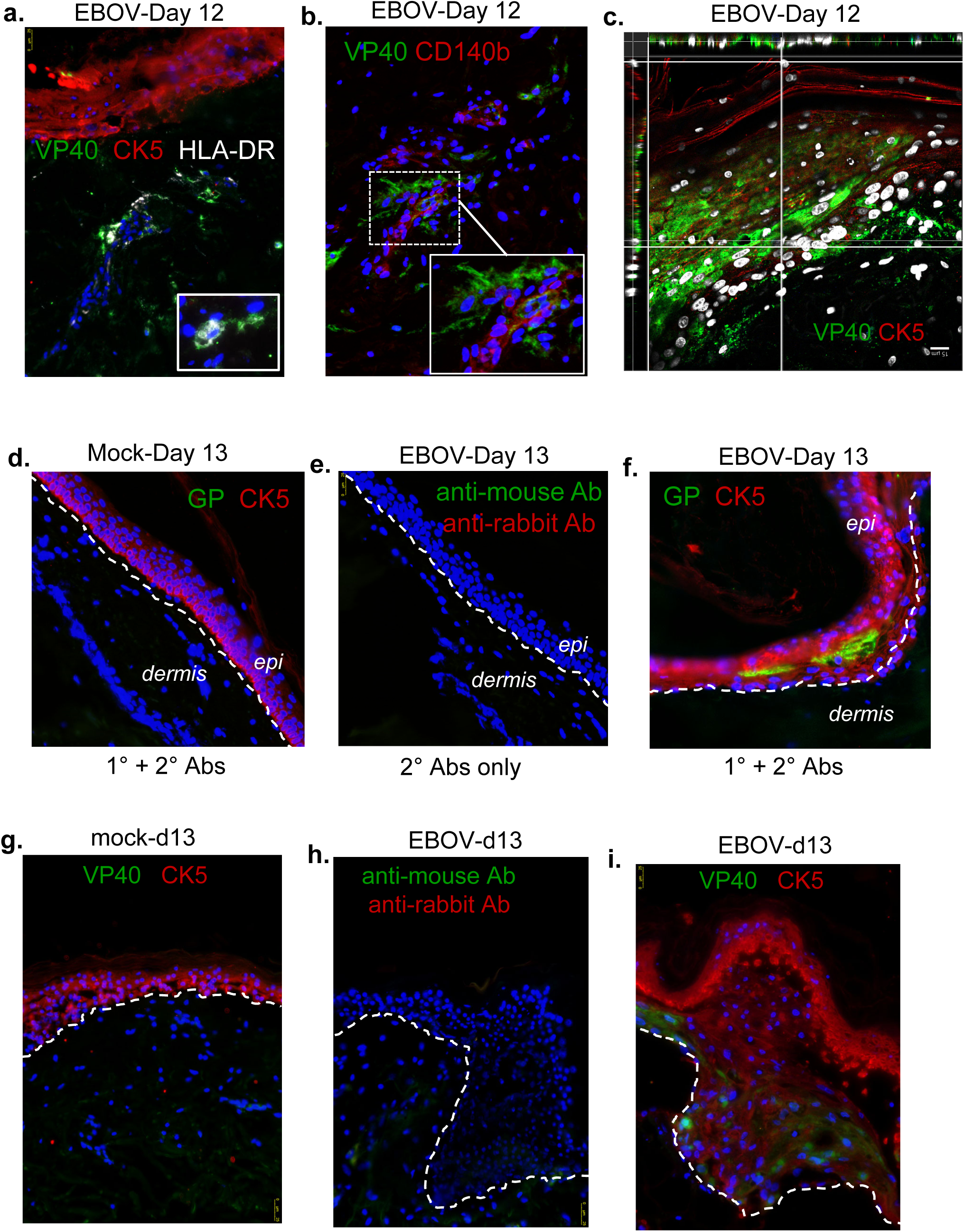
Multiple skin cell subsets support EBOV infection: additional evidence of BOV infection of skin explants. Human skin explants were generated and maintained as scribed in Figure 1. Explants were inoculated with 10^6^ FFU EBOV and harvested on days 8-13. Sections of FFPE explants were immunostained with antibodies for EBOV GP or VP40 (green), lineage-specific markers (red), and mounted with DAPI (blue/white). Lineage-specific markers were CK5 (epidermis), CD140b (fibroblast/smooth muscle cells/pericytes), CD31 (endothelia), and HLA-DR in white (antigen-presenting cell). The dashed line indicates the junction between the epidermis (epi) and the dermis. **a**) In explants harvested on d 12, VP40 colocalized with HLA-DR+ cells in the dermis, and (**b**) clusters of VP40+ cells surround a cluster of CD140b+ fibroblasts in the dermis. (**c**) An orthogonal z-stack projection shows intense viral staining that colocalizes with layers of CK5+ epidermal keratinocytes. Findings are representative studies of 3-5 independent experiments. For all staining experiments, controls include (**d, g**) specific staining of mock-infected tissue with both primary and secondary antibodies and (**e, h**) staining of virally infected explants with secondary (2°) antibodies alone, which are assessed to confirm specific staining of virally infected tissue (**f, i**). Scale bar = 15 μM in panel c and 25 μM in all others.

**Suppl. Fig. 3.**
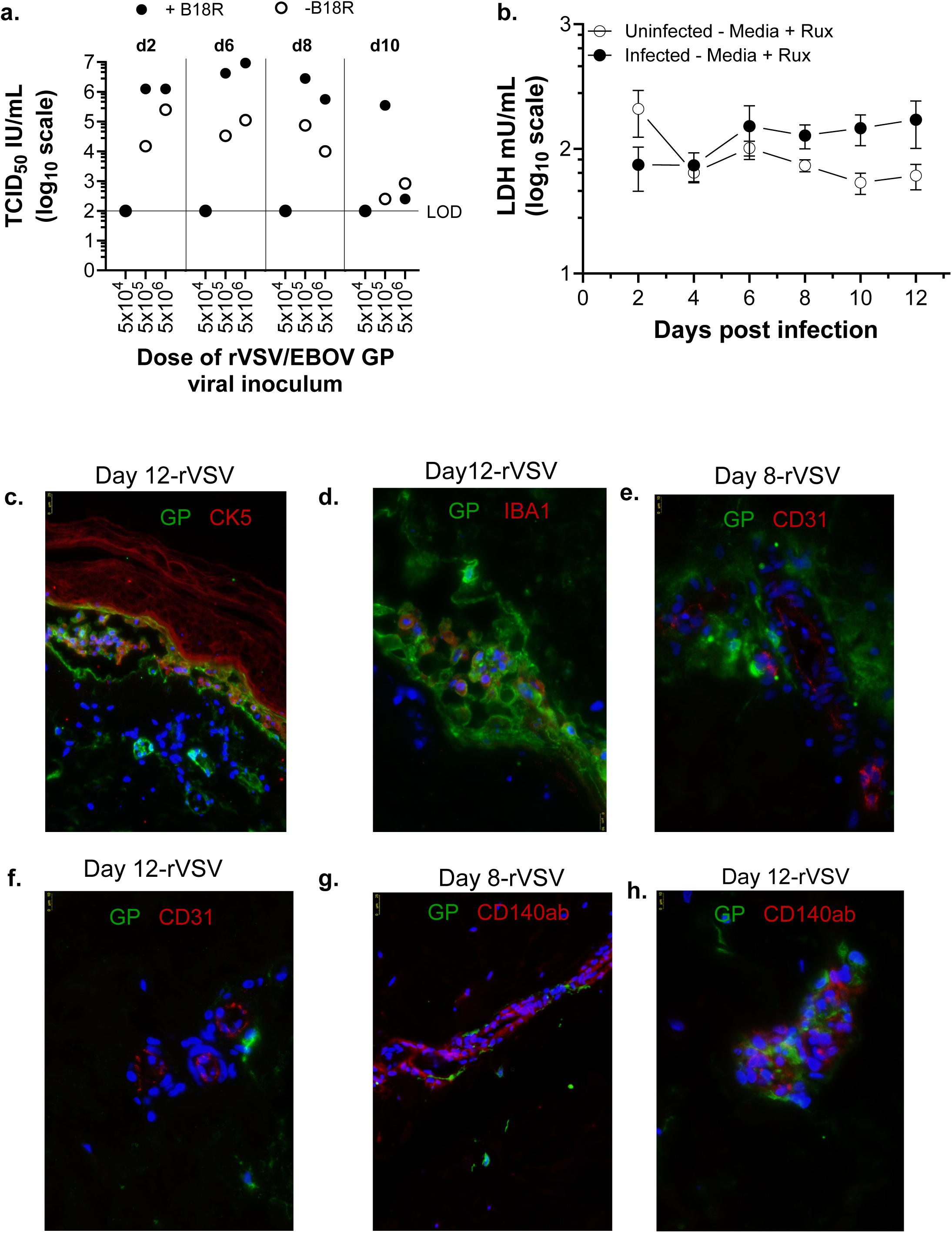
rVSV/EBOV tropism is similar to EBOV in human skin explants: additional evidence of robust infection. **a)** Titers of rVSV/EBOV GP in basal supernatants of foreskin explants over the course of a 10-day infection in the presence or absence of B18R. Explants were infected in the basal media with the dose of virus noted. Media was collected on the days noted and refreshed. Titers of virus were assessed by TCID_50_ assays on Vero cells. **b)** LDH was measured in basal supernatants of uninfected (open circles) and rVSV/EBOV GP-infected skin explants (black circles) cultured for 12 days with media changes every other day. Media for both conditions contained rux. **c-h)** FFPE explants were sectioned and immunostained with antibodies for EBOV GP (green) or lineage-specific antibodies (red) and mounted with DAPI (blue). Lineage markers included CK5 (epidermis), CD140b (fibroblast/smooth muscle cells/pericytes), CD31 (endothelia), and IBA1 (monocyte/macrophage). Scale bars = 10 μM in d, e, f, h and 25 μM in c, g.

**Suppl. Fig. 4.**
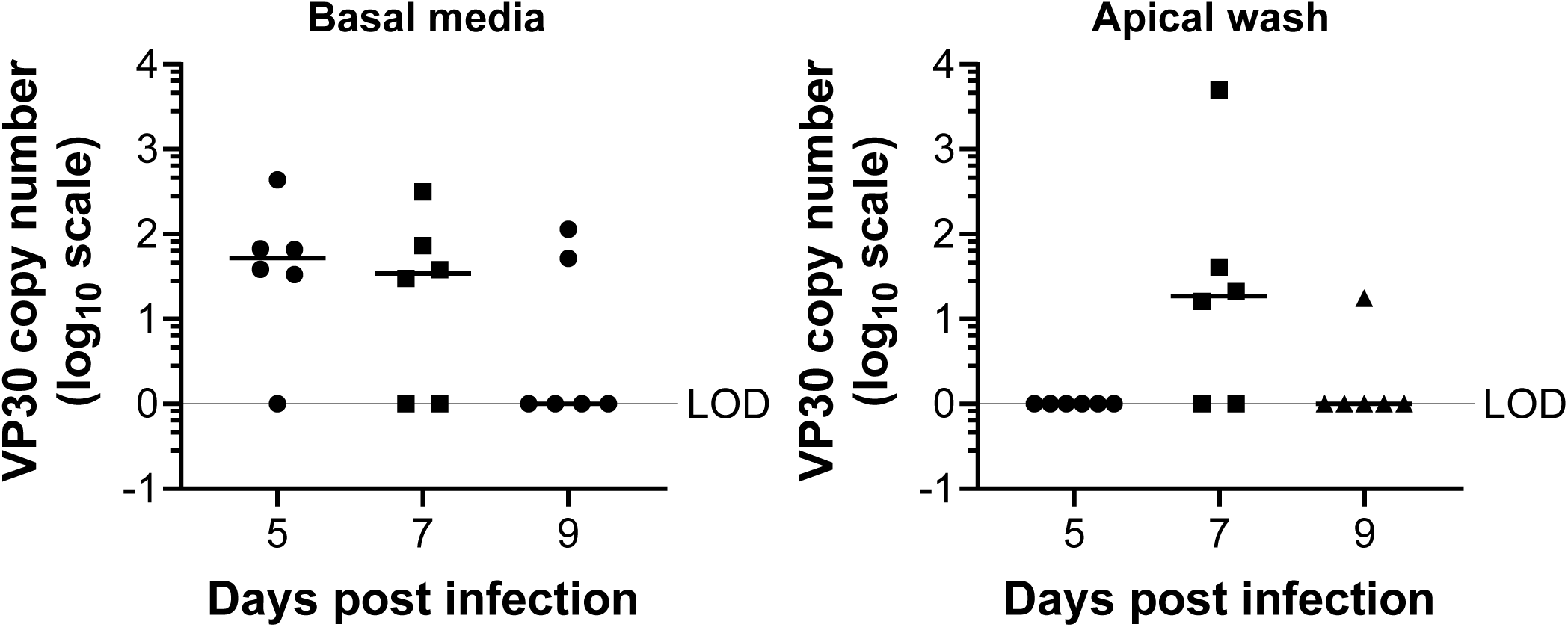
Virus traffics through the explants over time: detection of EBOV RNA in supernatants and the apical surface of explants. Detection of viral load in basal media (left panel) and apical PBS wash (right panel) from days 5-9 of EBOV-GFP infection. Explants were infected with 10^5^ of EBOV and input virus was removed after 18 hours. Media was collected and refreshed every other day. At the time of collection, basal supernatants were collected. Ten μl of sterile PBS was placed on the surface of the stratum corneum for 1 minute and this was repeated. Collected PBS was pooled and RNA extracted. Viral load was determined by assessing VP30 by qRT-PCR and the values were normalized to a VP30 standard curve to determine copy number.

**Suppl. Fig. 5.**
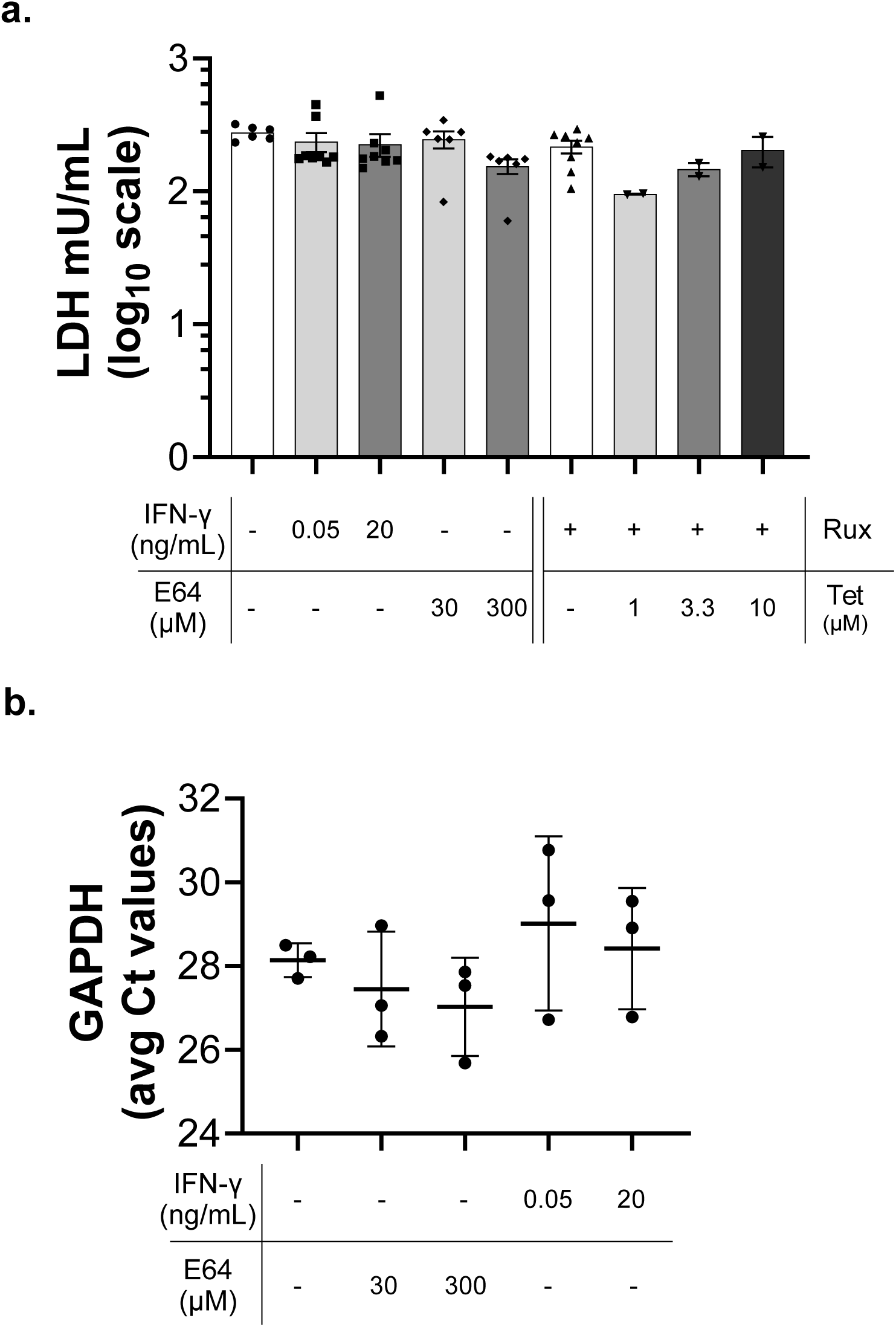
Human skin explants serve as excellent models for drug discovery: effect of drugs on cell viability. **a**) LDH levels detected in supernatant from day 12 uninfected explants that were maintained with the appropriate drug concentration in a transwell format with complete media changes every other day. Shown are means ± SEM. (n=2 independent donors). **b**) Expression of housekeeping gene, GAPDH, in uninfected day 14 explants that were maintained with the appropriate drug concentration in a transwell format with complete media changes every other day. Shown are means ± SD of three explants/treatment.

**Suppl. Fig.6.**
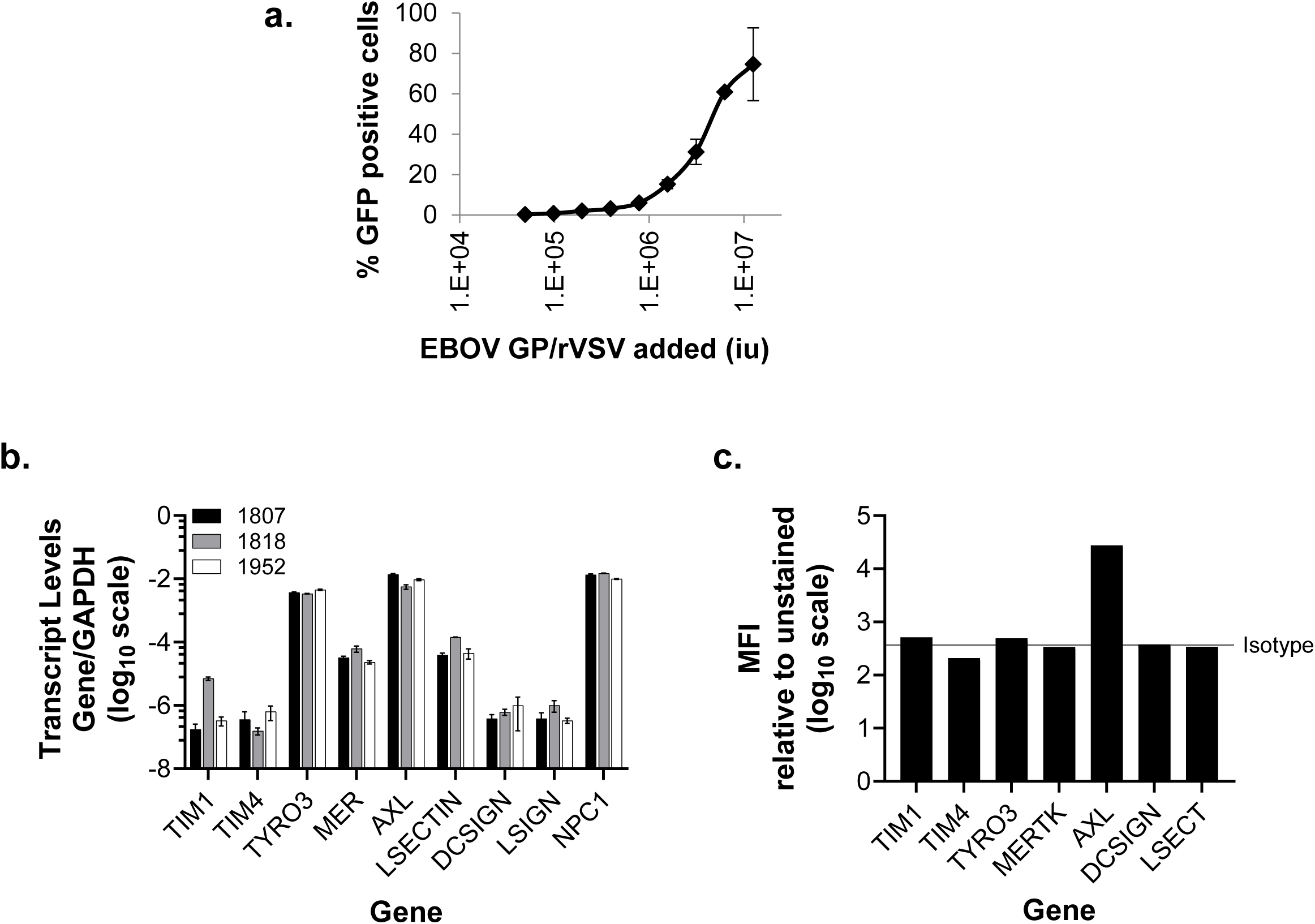
Primary and immortalized human keratinocytes support EBOV infection in an Axl- and NPC1-dependent manner: additional datasets. **a**) rVSV/EBOV GP dose response curve in primary keratinocytes. Shown is percent GFP positive cells at 24 hpi as measured by flow cytometry. Data are shown as mean and SD. (n=3 independent times). **b**) Gene expression of cell surface or endosomal receptors known to be used by filoviruses. **c**) Cell surface expression of receptors as assessed by flow cytometry on primary human keratinocytes.

**Suppl. Fig. 7.**
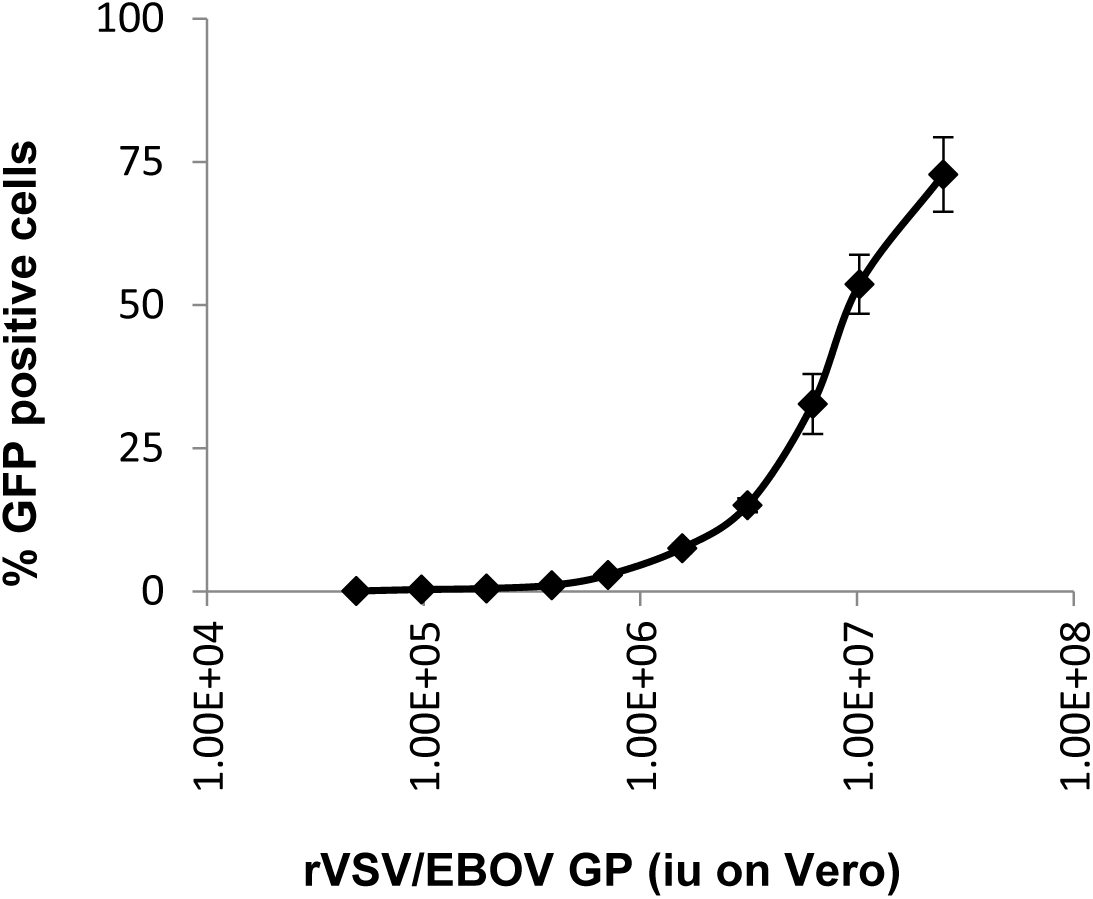
Primary and immortalized human skin fibroblasts support EBOV infection in an Axl- and NPC1-dependent manner: dose response curve of rVSV/EBOV GP on primary human skin fibroblasts. Cells were plated in a 48-well format and infected with the concentrations of rVSV/EBOV GP noted at confluency. At 24 hpi, cells were trypsinized and analyzed by flow cytometry for GFP expression. Shown are mean and SE. (n=3 independent experiments).

## Supplemental Movie Legends

**Supplemental movie 1. Localization of epidermal of EBOV VP40 and CK5 staining of skin explants.** Explants were inoculated with 10^6^ FFU EBOV and sections from explants harvested on d12 post-infection were immunostained with antibodies for EBOV (green), CD5 (red), and counter-stained with DAPI (blue). In this image, the section is oriented vertically with the CK5^+^ epidermis on left and the dermis on right. Viral staining colocalizes with CK5 staining in the basal and suprabasal layers of the epidermis. Staining was visualized on z-series image stacks using high-resolution confocal microscopy. Scale bar =25μM.

**Supplemental movie 2. Localization of dermal rVSV/EBOV GP and CD140ab staining of skin explants.** Explants were inoculated with 10^7^ rVSV and sections from explants harvested on d8 post-infection were immunostained with antibodies for EBOV GP (green), CD140ab (red), and DAPI (blue). CD140ab staining is primarily evident in the thicker layers of a dermal arteriole, surrounded by a thin layer viral GP stain, consistent with fibroblasts or pericytes. In the lower left quadrant, viral GP staining is observed in a large single cell with morphology consistent with a macrophage. Staining was visualized on z-series image stacks using high-resolution confocal microscopy. Scale bar =25μM.

**Supplemental movie 3. Localization of dermal of rVSV/EBOV GP and CD31 staining of skin explants.** Explants were inoculated with 10^7^ rVSV and sections from explants harvested on d8 post-infection were immunostained with antibodies for EBOV GP (green), CD31 (red), and DAPI (blue). CD31 staining is evident in the endothelial layers of dermal vessels surrounded by cellular and diffuse viral GP staining. The absence of colocalization suggests that endothelial cells are not infected in these samples. Staining was visualized on z-series image stacks using high-resolution confocal microscopy. Scale bar =25μM.

