## Supplementary material for "Multiple cell types support productive infection and dynamic translocation of infectious Ebola virus to the surface of human skin": Suppl. Fig. 1

**Figure S1**

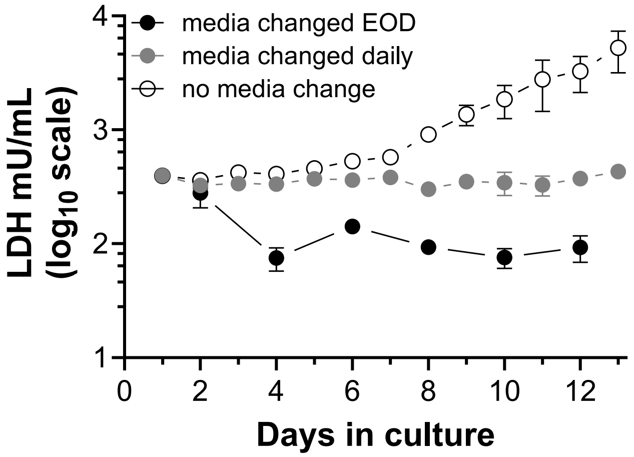


**Suppl. Fig. 1. Human skin explant model of EBOV infection: LDH levels in supernatant of uninfected human skin explants under different culture conditions.** As an index of viability, lactate dehydrogenase activity (LDH) was measured in supernatants of uninfected skin explants cultured for 13 days without any media change (open circles), with complete media change every day (grey circles), or every other day (black circles). Data are expressed as mean ± SD. n = 6 replicates per condition from a single donor.

**Figure S2**

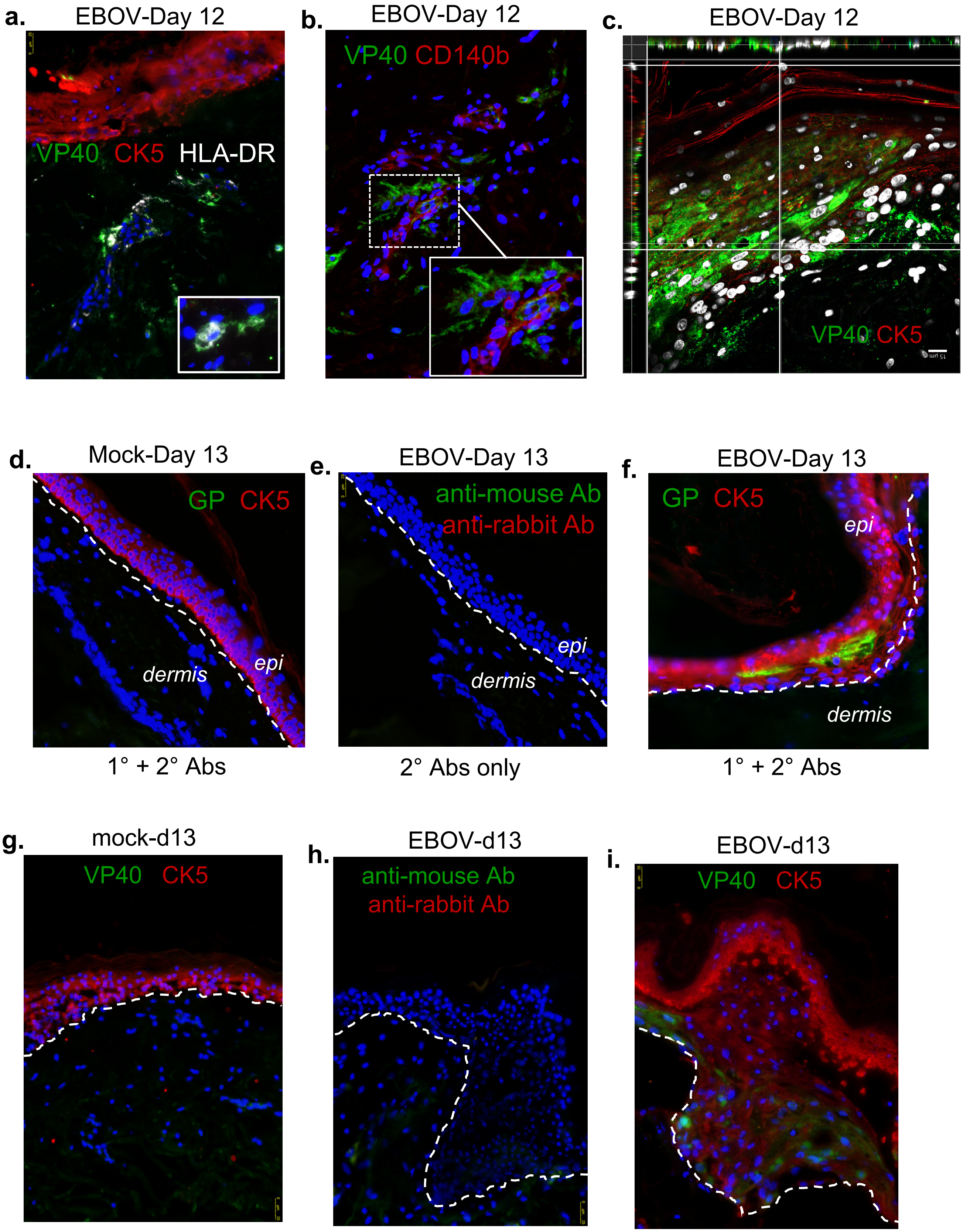

a.
EBOV-Day 12
VP40 CK5 HLA-DR
b.
EBOV-Day 12
VP40 CD140b
c.
EBOV-Day 12
VP40 CK5
d.
Mock-Day 13
GP CK5
dermis
epi
1° + 2° Abs
e.
EBOV-Day 13
anti-mouse Ab
anti-rabbit Ab
epi
dermis
2° Abs only
f.
EBOV-Day 13
GP CK5
epi
dermis
1° + 2° Abs
g.
mock-d13
VP40 CK5
h.
EBOV-d13
anti-mouse Ab
anti-rabbit Ab
i.
EBOV-d13
VP40 CK5

**Suppl. Fig. 2: Multiple skin cell subsets support EBOV infection: additional evidence of EBOV infection of skin explants.** Human skin explants were generated and maintained as described in Figure 1. Explants were inoculated with $10^6$ FFU EBOV and harvested on days 8-13. Sections of FFPE explants were immunostained with antibodies for EBOV GP or VP40 (green), lineage-specific markers (red), and mounted with DAPI (blue/white). Lineage-specific markers were CK5 (epidermis), CD140b (fibroblast/smooth muscle cells/pericytes), CD31 (endothelia), and HLA-DR in white (antigen-presenting cell). The dashed line indicates the junction between the epidermis (epi) and the dermis. **a**) In explants harvested on d 12, VP40 colocalized with HLA-DR+ cells in the dermis, and (**b**) clusters of VP40+ cells surround a cluster of CD140b+ fibroblasts in the dermis. (**c**) An orthogonal z-stack projection shows intense viral staining that colocalizes with layers of CK5+ epidermal keratinocytes. Findings are representative studies of 3-5 independent experiments. For all staining experiments, controls include (**d, g**) specific staining of mock-infected tissue with both primary and secondary antibodies and (**e, h**) staining of virally infected explants with secondary (2°) antibodies alone, which are assessed to confirm specific staining of virally infected tissue (**f, i**)**.** Scale bar = 15 μM in panel c and 25 μM in all others.

### Figure S3

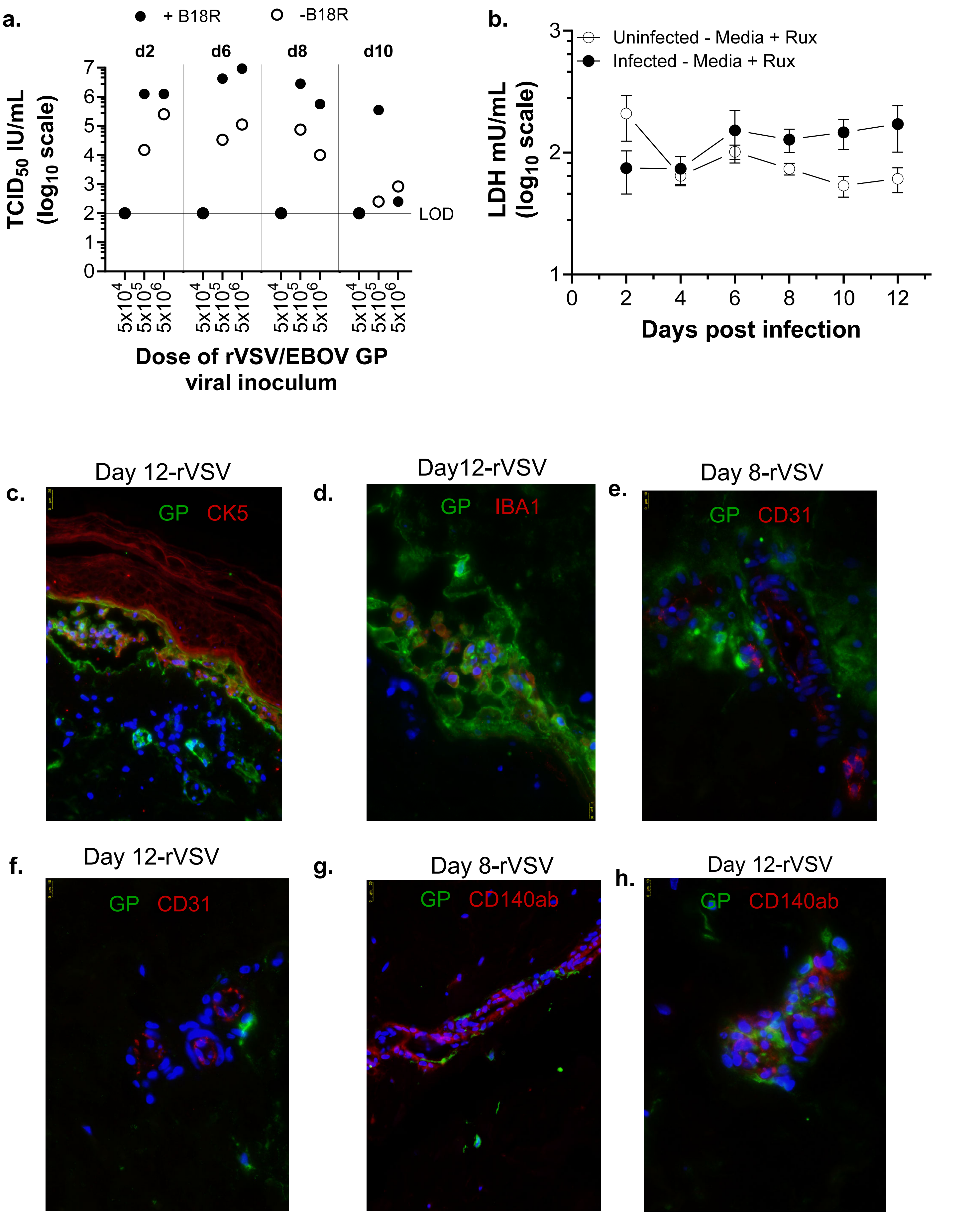


Suppl. Fig. 3

**Suppl. Fig. 3. rVSV/EBOV tropism is similar to EBOV in human skin explants: additional evidence of robust infection. a)** Titers of rVSV/EBOV GP in basal supernatants of foreskin explants over the course of a 10-day infection in the presence or absence of B18R. Explants were infected in the basal media with the dose of virus noted. Media was collected on the days noted and refreshed. Titers of virus were assessed by $TCID_{50}$ assays on Vero cells. **b)** LDH was measured in basal supernatants of uninfected (open circles) and rVSV/EBOV GP-infected skin explants (black circles) cultured for 12 days with media changes every other day. Media for both conditions contained rux. **c-h**) FFPE explants were sectioned and immunostained with antibodies for EBOV GP (green) or lineage-specific antibodies (red) and mounted with DAPI (blue). Lineage markers included CK5 (epidermis), CD140b (fibroblast/smooth muscle cells/pericytes), CD31 (endothelia), and IBA1 (monocyte/macrophage). Scale bars = 10 µM in d, e, f, h and 25 µM in c, g.

**Figure S4**

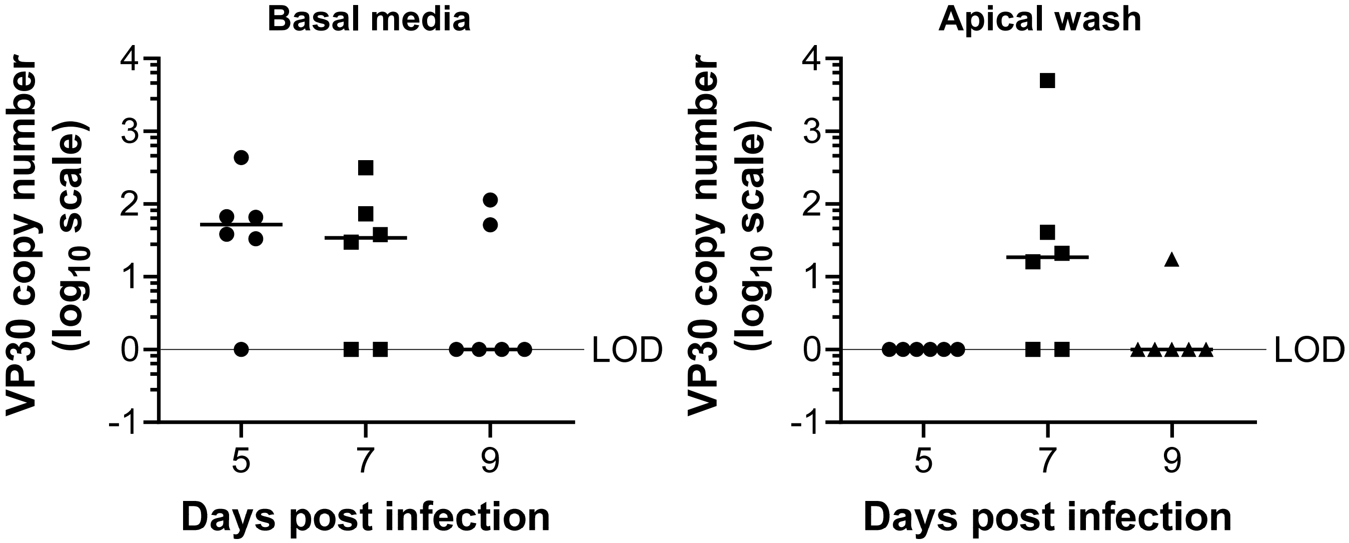


**Suppl. Fig. 4. Virus traffics through the explants over time: detection of EBOV RNA in supernatants and the apical surface of explants.** Detection of viral load in basal media (left panel) and apical PBS wash (right panel) from days 5-9 of EBOV-GFP infection. Explants were infected with $10^5$ of EBOV and input virus was removed after 18 hours. Media was collected and refreshed every other day. At the time of collection, basal supernatants were collected. Ten µl of sterile PBS was placed on the surface of the stratum corneum for 1 minute and this was repeated. Collected PBS was pooled and RNA extracted. Viral load was determined by assessing VP30 by qRT-PCR and the values were normalized to a VP30 standard curve to determine copy number.

### Figure S5

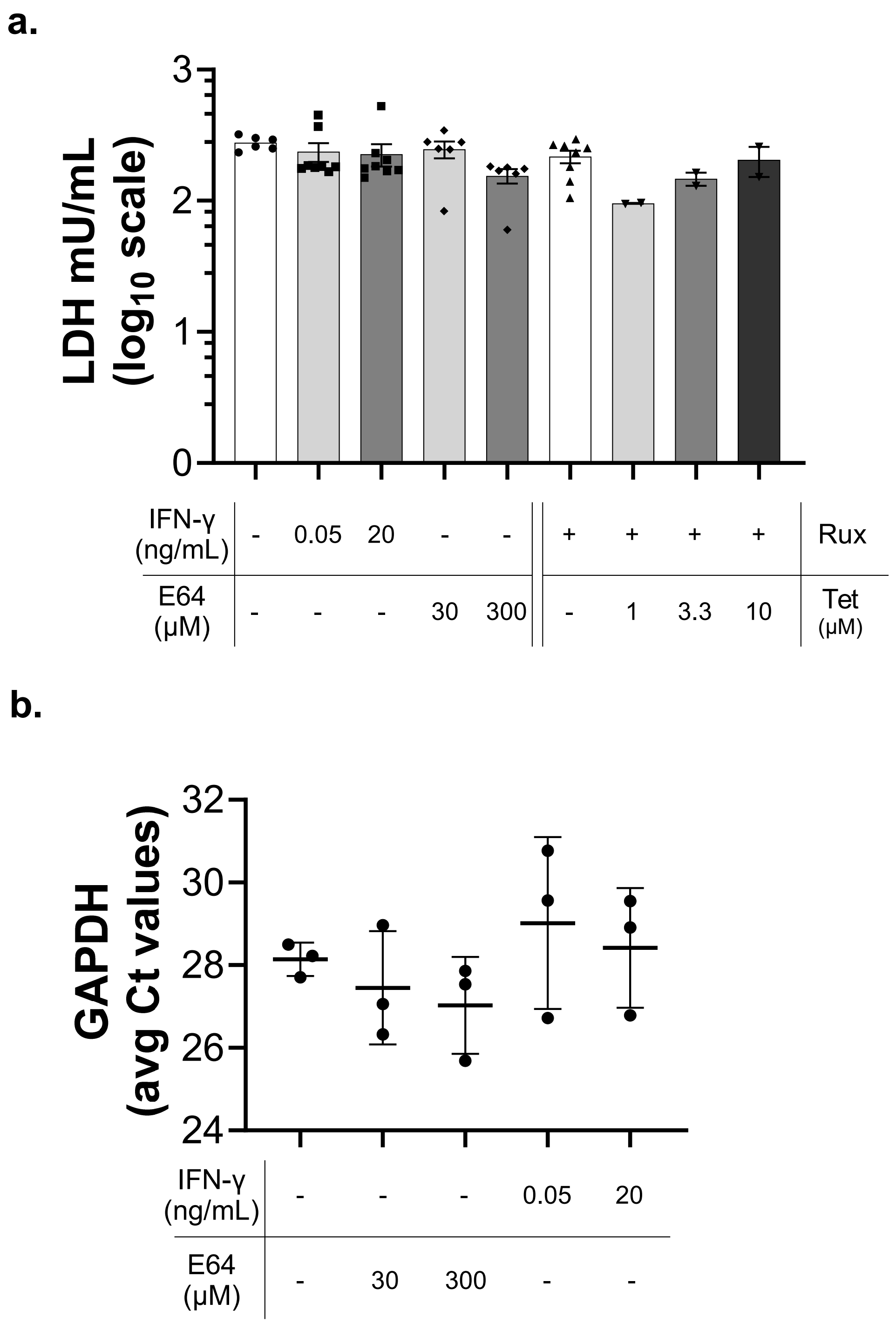


**Suppl. Fig. 5**. **Human skin explants serve as excellent models for drug discovery: effect of drugs on cell viability. a**) LDH levels detected in supernatant from day 12 uninfected explants that were maintained with the appropriate drug concentration in a transwell format with complete media changes every other day. Shown are means ± SEM. (n=2 independent donors). **b**) Expression of housekeeping gene, GAPDH, in uninfected day 14 explants that were maintained with the appropriate drug concentration in a transwell format with complete media changes every other day. Shown are means ± SD of three explants/treatment.

**Figure S6**

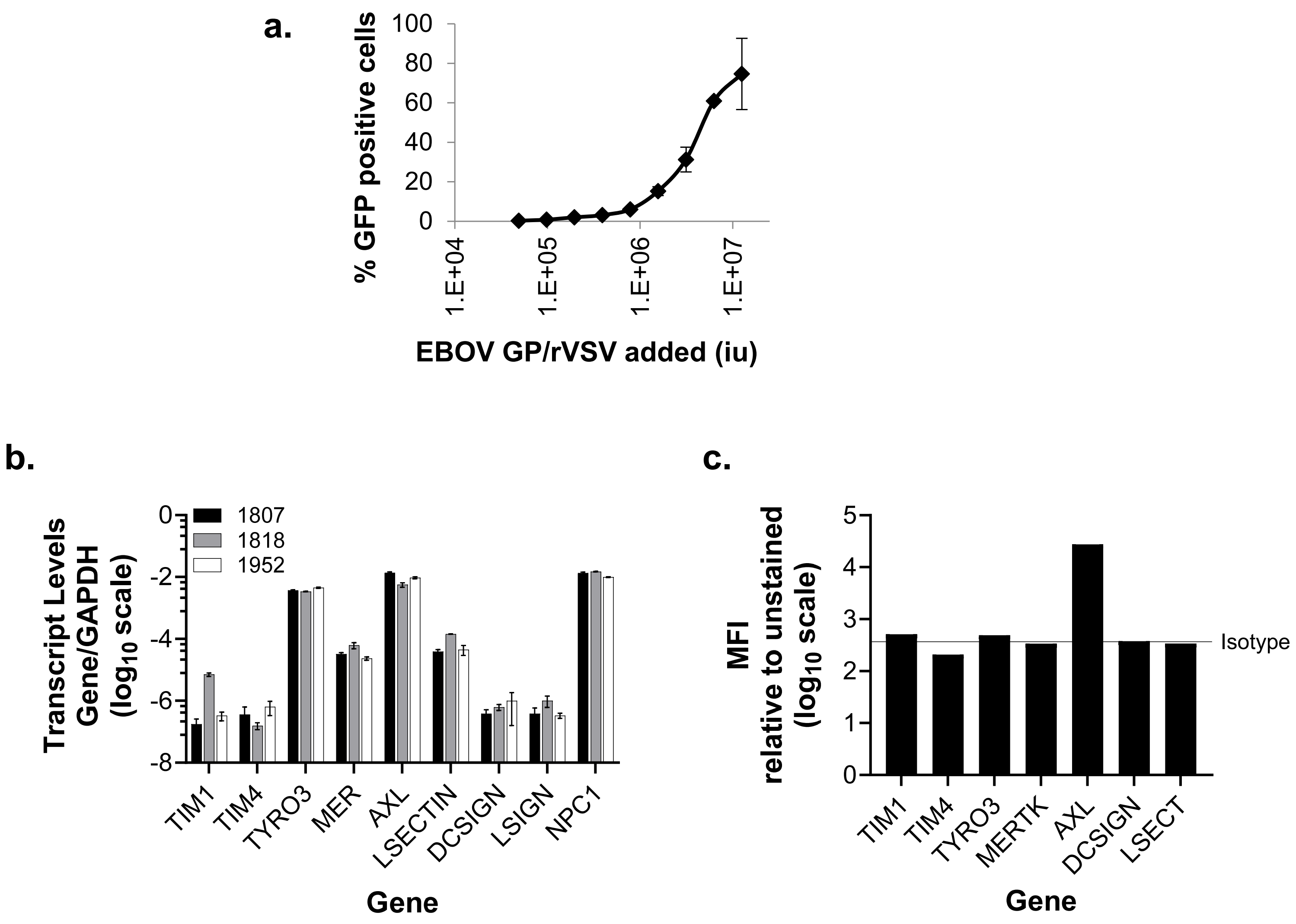


**Suppl. Fig.6. Primary and immortalized human keratinocytes support EBOV infection in an Axl- and NPC1-dependent manner: additional datasets. a**) rVSV/EBOV GP dose response curve in primary keratinocytes. Shown is percent GFP positive cells at 24 hpi as measured by flow cytometry. Data are shown as mean and SD. (n=3 independent times). **b**) Gene expression of cell surface or endosomal receptors known to be used by filoviruses. **c**) Cell surface expression of receptors as assessed by flow cytometry on primary human keratinocytes.

#### Figure S7

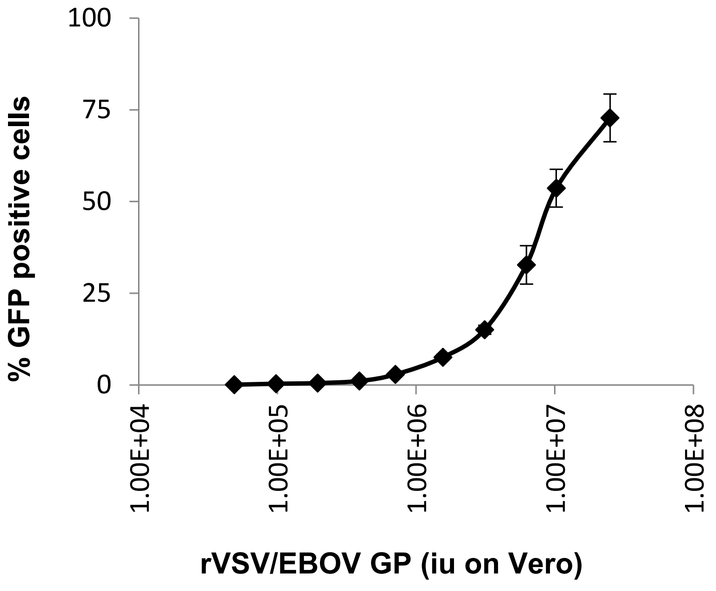


**Suppl. Fig. 7. Primary and immortalized human skin fibroblasts support EBOV infection in an Axl- and NPC1-dependent manner: dose response curve of rVSV/EBOV GP on primary human skin fibroblasts.** Cells were plated in a 48-well format and infected with the concentrations of rVSV/EBOV GP noted at confluency. At 24 hpi, cells were trypsinized and analyzed by flow cytometry for GFP expression. Shown are mean and SE. (n=3 independent experiments).

**Supplemental Movie Legends:**

**Supplemental movie 1. Localization of epidermal of EBOV VP40 and CK5 staining of skin explants.** Explants were inoculated with $10^6$ FFU EBOV and sections from explants harvested on d12 post-infection were immunostained with antibodies for EBOV (green), CD5 (red), and counter-stained with DAPI (blue). In this image, the section is oriented vertically with the $CK5^+$ epidermis on left and the dermis on right. Viral staining colocalizes with CK5 staining in the basal and suprabasal layers of the epidermis. Staining was visualized on z-series image stacks using high-resolution confocal microscopy. Scale bar =25µM.

**Supplemental movie 2. Localization of dermal rVSV/EBOV GP and CD140ab staining of skin explants.** Explants were inoculated with $10^7$ rVSV and sections from explants harvested on d8 post-infection were immunostained with antibodies for EBOV GP (green), CD140ab (red), and DAPI (blue). CD140ab staining is primarily evident in the thicker layers of a dermal arteriole, surrounded by a thin layer viral GP stain, consistent with fibroblasts or pericytes. In the lower left quadrant, viral GP staining is observed in a large single cell with morphology consistent with a macrophage. Staining was visualized on z-series image stacks using high-resolution confocal microscopy. Scale bar =25µM.

**Supplemental movie 3. Localization of dermal of rVSV/EBOV GP and CD31 staining of skin explants.** Explants were inoculated with $10^7$ rVSV and sections from explants harvested on d8 post-infection were immunostained with antibodies for EBOV GP (green), CD31 (red), and DAPI (blue). CD31 staining is evident in the endothelial layers of dermal vessels surrounded by cellular and diffuse viral GP staining. The absence of colocalization suggests that endothelial cells are not infected in these samples. Staining was visualized on z-series image stacks using high-resolution confocal microscopy. Scale bar =25µM.
